# Multi-stromal organoid co-cultures model pancreatic cancer and pancreatitis epithelial cell-fibroblast heterogeneity

**DOI:** 10.64898/2025.12.05.692494

**Authors:** Wenlong Li, Muntadher Jihad, Eloise G. Lloyd, Gianluca Mucciolo, Joaquín Araos Henríquez, Marta Zaccaria, Priscilla S.W. Cheng, Sneha Harish, Sally Mills, Paul M. Johnson, Weike Luo, Alejandro Alonso Montero, Alex Deamer, Rebecca Brais, Ania M. Piskorz, Paul D.W. Kirk, Mireia Vallespinos, Giulia Biffi

## Abstract

Malignant cell-fibroblast cross-talks modulate disease progression and therapy response of pancreatic ductal adenocarcinoma (PDAC). Our knowledge of the heterogeneous nature of PDAC cancer-associated fibroblasts (CAFs) has significantly increased over the last few years. Yet, whether CAFs in PDAC differ from fibroblasts in pancreatic inflammation remains poorly understood. Chronic pancreatitis – a prolonged inflammatory state of the pancreas – is a risk factor for PDAC and is characterised by abundant fibroblasts. Thus, dissecting pancreatic fibroblast and epithelial cell reprogramming in malignancy relative to inflammation could inform new preventative, diagnostic and therapeutic strategies for PDAC. Here, we studied how pancreatic malignancy and inflammation differently shape fibroblast heterogeneity and their crosstalk with epithelial cells. We analysed human samples and mouse models of pancreatitis and PDAC and leveraged new murine pancreatitis-derived epithelial organoids to establish pancreatitis and PDAC organoid/multi-stroma co-cultures comprising pancreatic stellate cells, fibroblasts and mesothelial cells. We demonstrate that a combination of *in vitro* and *in vivo* models better captures epithelial cell and fibroblast markers of human pancreatitis and PDAC compared to mouse models alone. Finally, we identify PDAC and pancreatitis epithelial cell-specific reprogramming of stromal cells of different origin, and we infer the contribution of these distinct stromal cell types to fibroblasts in PDAC and pancreatitis *in vivo*. Together, our study highlights different epithelial cell-fibroblast heterogeneity in PDAC and pancreatitis, and provides new platforms for the identification of markers and epithelial-stromal interdependencies of these diseases.

## INTRODUCTION

The high mortality rate of pancreatic ductal adenocarcinoma (PDAC) is largely attributed to the late-stage of diagnosis, treatment resistance, and extensive non-cancerous stroma^1^. Within this stroma, cancer-associated fibroblasts (CAFs) are known determinants of PDAC progression and therapy response^2,3^. PDAC CAFs are phenotypically and functionally heterogenous, typically classified into three states: inflammatory CAFs (iCAFs), myofibroblastic CAFs (myCAFs) and antigen-presenting CAFs (apCAFs). However, additional populations within and across these states have also been described^4–8^. Whether fibroblasts in PDAC are similar to those in non-malignant pancreatic diseases, is less clear. Comparing CAFs in PDAC to fibroblasts in pancreatic conditions that are risk factors for this malignancy could help identify cancer-specific fibroblast states that may be important for the development of preventative, diagnostic and therapeutic PDAC strategies.

In humans, chronic pancreatitis is a persistent inflammatory state and a risk factor of PDAC^9,10^. It is characterised by increased fibrosis and extracellular matrix (ECM) deposition that contribute to its progression^11^. The ECM of chronic pancreatitis shares features with that in PDAC both in murine models and human tissues^12^. This may be due to the fact that activated, ECM-producing pancreatic stellate cells (PSCs) are, at least in part, precursors of PDAC CAFs as well as contributors to the fibrosis of pancreatitis^13–15^.

Mouse models have advanced our understanding of how epithelial cells are reprogrammed during pancreatic injury and inflammation, and how this contributes to malignant transformation^16–19^. Yet, less is known about how fibroblasts are reprogrammed during pancreatitis. Indeed, while previous works showed that fibroblasts in pancreatitis are heterogeneous^15,20^, potential differences in fibroblasts, and epithelial cell-fibroblast crosstalk, between pancreatitis and PDAC remain underexplored. This is partially due to the lack of pancreatitis models that enable the dissection of epithelial cell-fibroblast crosstalk. The most commonly deployed mouse model of pancreatitis involves caerulein-induced pancreatic injury, which promotes release of digestive enzymes from acinar cells, acinar-to-ductal metaplasia (ADM) and fibrosis^21^. The establishment of additional models of pancreatitis may enable orthogonal validation of findings in mouse models, as well as capture a broader range of human pancreatitis- and PDAC- specific features.

Here, we aimed to identify epithelial cell and fibroblast features that are characteristic of pancreatic inflammation or malignancy. To this end, we analysed murine and human pancreatitis and PDAC tissues, as well as new pancreatitis- and PDAC-derived organoid/multi-stroma co-culture models.

## RESULTS

### Myofibroblasts are more abundant in human and murine PDAC relative to pancreatitis

To evaluate whether epithelial cells and fibroblasts have distinct features in PDAC compared to pancreatitis, we first analysed a published single-cell RNA-sequencing (scRNA-seq) dataset of human PDAC (n=5) and chronic pancreatitis (n=4; **Fig. 1A-C** and **Extended Data Fig. 1A-C** and **Supplementary Table 1**)^20^. Proliferation-associated (e.g., G2M checkpoint) and inflammatory (e.g., inflammatory response) pathways were upregulated in PDAC malignant cells compared to pancreatitis ductal cells (**Fig. 1D** and **Supplementary Table 1**). We also found an upregulation of myofibroblastic signatures (e.g., smooth muscle contraction, collagen formation) in PDAC CAFs compared to pancreatitis fibroblasts (**Fig. 1E** and **Supplementary Table 1**). These findings indicate that epithelial and fibroblast transcriptomes differ markedly between human pancreatitis and PDAC.

**Figure 1.**
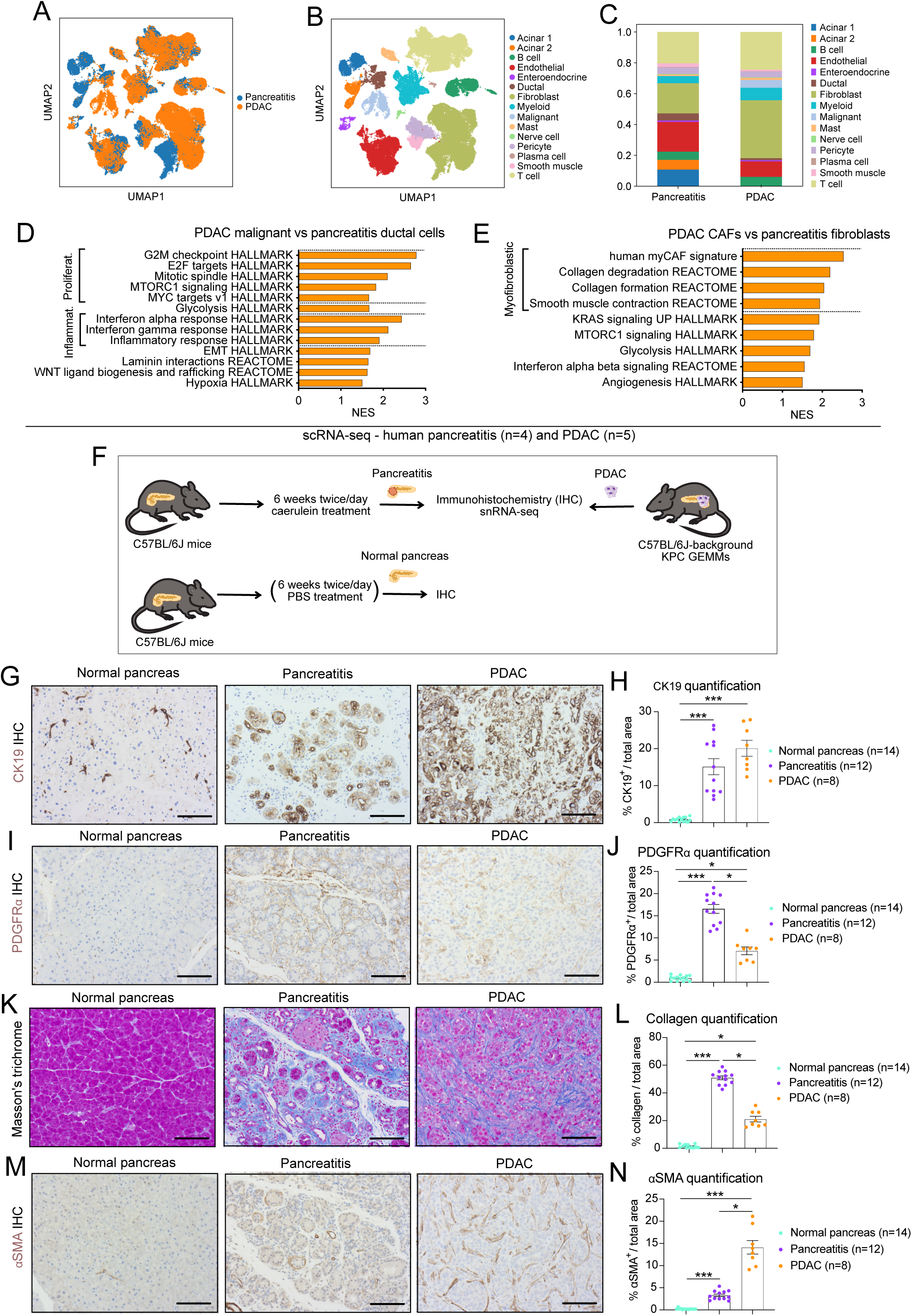
Myofibroblasts are more abundant in human and murine PDAC relative to pancreatitis. **(A-B)** Uniform Manifold Approximation and Projection (UMAP) plots of cell types in the combined single-cell RNA-sequencing (scRNA-seq) dataset from human pancreatic ductal adenocarcinoma (PDAC) (n=5) and chronic pancreatitis (n=4) tissues. Different conditions **(A)** and cell types **(B)** are colour coded. scRNA-seq data is from Dimitrieva et al. **(C)** Cell type contribution in human PDAC and pancreatitis, represented as bar plot showing proportions of the different cell clusters in each condition. **(D)** Significantly upregulated pathways identified by gene set enrichment analysis (GSEA) of PDAC malignant cells compared to pancreatitis ductal cells, as assessed by Model-based Analysis of Single-cell Transcriptomics (MAST) from the scRNA-seq dataset, and genes were ranked based on log2 fold change. Inflammat, inflammatory; Proliferat., proliferative. **(E)** Significantly upregulated pathways identified by GSEA of PDAC cancer-associated fibroblasts (CAFs) compared to pancreatitis fibroblasts, as assessed by MAST from the scRNA-seq dataset. **(F)** Schematic of models and techniques used for analysis of normal pancreas (i.e. from untreated or 6-week PBS-treated mice), pancreatitis (i.e. from 6-week caerulein-treated mice) and PDAC (i.e. from KPC genetically engineered mouse models, GEMMs) tissues from C57BL/6J mice. **(G-N)** Representative images and quantification of the ductal marker keratin 19 (CK19) **(G-H)**, the fibroblast marker PDGF receptor alpha (PDGFRα) **(I-J)**, Masson’s trichrome for collagen deposition **(K-L)** and the myofibroblast marker alpha smooth muscle actin (αSMA) **(M-N)** immunohistochemical stains of normal pancreas, pancreatitis and PDAC tissues. Results show mean ± SEM. * *P adj* < 0.05; ** *P adj* < 0.01; *** *P adj* < 0.001, Kruskal-Wallis test. Scale bars, 50 μm.

To further understand how epithelial cells and fibroblasts differ between pancreatitis and PDAC, we established mouse models of pancreatitis and compared tissues from these models to murine normal pancreas and PDAC tissues. PDACs were obtained from KPC (*Kras*^LSL-G12D/+^; *Trp53*^LSL-R172H/+^; *Pdx1*-Cre) genetically engineered mouse models (GEMMs)^22^, while pancreatitis tissues were obtained from C57BL/6J mice treated with the cholecystokinin analogue caerulein^21^ (**Fig. 1F** and **Extended Data Fig. 1D**). We established models of pancreatitis and acute pancreatitis by treating mice with caerulein for six weeks or two days, respectively (**Fig. 1F** and **Extended Data Fig. 1D**). Normal pancreata were obtained from untreated C57BL/6J mice or these mice injected with phosphate-buffered saline (PBS, vehicle) for six-weeks (i.e., normal mice; **Fig. 1F**). As expected, caerulein treatment reduced plasma amylase and lipase activity levels and pancreas weights in acute pancreatitis and pancreatitis mice compared to normal mice, compatible with acinar damage and pancreatic atrophy (**Extended Data Fig. 1E-G**)^23^. Pancreas weights were reduced the most in pancreatitis mice (**Extended Data Fig. 1G**).

Pancreatitis is associated with loss of acinar cells through ADM, increased immune cell infiltration and fibrosis^24^. Therefore, we used immunohistochemistry to compare these features among pancreatitis, acute pancreatitis and normal pancreas tissues. Expression of the ductal marker cytokeratin 19 (CK19) was significantly increased in pancreatitis and acute pancreatitis compared to normal pancreas tissues, indicative of elevated ADM (**Fig. 1G-H** and **Extended Data Fig. 1H-I**). Pancreatitis and acute pancreatitis also displayed significantly increased levels of immune cell infiltration compared to normal pancreas tissues, as indicated by increased expression of F4/80 and Ly6G, which mark macrophages and neutrophils, respectively (**Extended Data Fig. 1H-M**). Finally, significantly elevated levels of fibrosis were also observed in pancreatitis and acute pancreatitis compared to normal pancreas tissues, which we detected using: fibroblast markers platelet-derived growth factor receptor alpha (PDGFRα) and podoplanin (PDPN), myofibroblast marker alpha smooth muscle actin (αSMA), and Masson’s trichrome stain for collagen (**Fig. 1I-N** and **Extended Data Fig. 1H-I** and **1N-O**). Pancreatitis tissues had higher levels of all these markers of fibrosis also relative to acute pancreatitis tissues (**Extended Data Fig. 1I**). These data are in line with evidence that prolonged inflammation regimens in mice better mimic the chronic inflammatory state associated with increased PDAC risk in patients^21^. Thus, we focused on pancreatitis, rather than acute pancreatitis, mouse models to further investigate how fibroblast and epithelial cells differ between pancreatic inflammation and PDAC.

To evaluate differences in stromal composition between murine pancreatitis and PDAC, we used our immunohistochemical strategy to compare pancreatitis and PDAC tissues. While levels of ADM (CK19) and macrophage infiltration (F4/80) were similar between conditions, neutrophil infiltration (Ly6G) was significantly greater in PDAC compared to pancreatitis (**Fig. 1G-H** and **Extended Data Fig. 1J-M**). We also noted differences in fibroblast composition. Specifically, levels of PDGFRα, PDPN and collagen were significantly higher in pancreatitis compared to PDAC tissues (**Fig. 1I-L** and **Extended Data Fig. 1N-O**). Conversely, myofibroblast abundance (αSMA) was higher in PDAC, compatible with our finding of increased myofibroblastic signatures in human PDAC CAFs relative to pancreatitis fibroblasts (**Fig. 1M-N and 1E**).

Together these data suggest that epithelial and stromal cells have distinct features in PDAC compared to pancreatitis tissues. Specifically, myofibroblasts are more abundant in murine and human PDAC.

### Fibroblast and epithelial cell populations are more heterogeneous in pancreatitis than PDAC mouse models

To further investigate differences between murine pancreatitis and PDAC stromal composition, we performed single-nuclei RNA-sequencing (snRNA-seq) of pancreatitis (n=6) and PDAC (n=4)^25^. The malignant and non-malignant cell populations in GEMM-derived PDACs resembled those previously found in murine organoid-derived PDACs^7^ (**Fig. 2A** and **Extended Data Fig. 2A-C**). Pancreatitis tissues contained additional cell populations that included four fibroblast subsets, which we termed ‘Fibro 1’ (*Wt1*^High^*Cd55*^High^), ‘Fibro 2’ (*Pdgfra*^High^*Pi16*^High^), ‘Fibro 3’ (*Fap*^High^) and ‘Fibro 4’ (*Wt1*^High^*Dpp4*^High^; **Fig. 2B-E** and **Extended Data Fig. 2D-G**). The *Fap*^High^ Fibro 3 subset was most similar to CAFs compared to other pancreatitis fibroblast subsets, as indicated by a greater enrichment of the PDAC CAF signature (i.e., top 50 marker genes; **Fig. 2F** and **Supplementary Table 2**). Furthermore, mapping the Fibro 1-4 signatures onto the PDAC CAF reference using a semi-supervised scANVI framework identified the highest transcriptional similarity between Fibro 3 and PDAC CAFs (31%; **Fig. 2G**)^26^. Pseudotemporal trajectory analysis further suggested that the Fibro 3 subset represents a direct precursor of PDAC CAFs (**Fig. 2H**).

**Figure 2.**
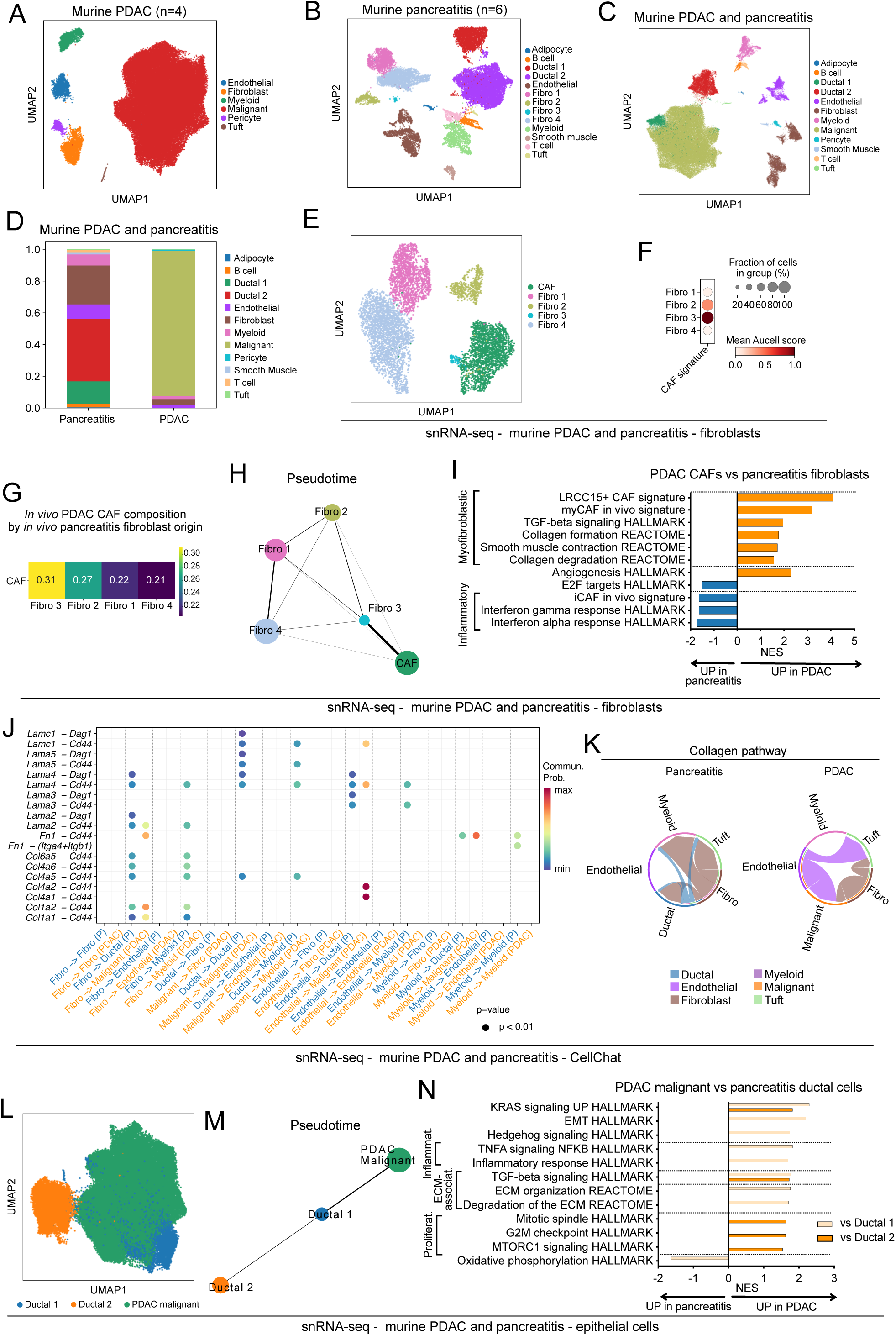
Fibroblast and epithelial cell populations are more heterogeneous in pancreatitis than PDAC mouse models. **(A-B)** UMAP plots of cell types in murine PDAC (n=4) **(A)** and pancreatitis (n=6) **(B)** tissues identified by single-nuclei RNA-sequencing (snRNA-seq). Different cell types are colour coded. **(C)** UMAP plot of cell types in the combined snRNA-seq dataset from murine PDAC and pancreatitis tissues. Different conditions cell types are colour coded. **(D)** Cell type contribution in murine PDAC and pancreatitis, represented as bar plot showing proportions of the different cell clusters in each condition. **(E)** UMAP plot of fibroblasts in the combined snRNA-seq dataset from murine PDAC and pancreatitis. Different fibroblast subsets are colour coded. **(F)** Dot plot of scaled Aucell score enrichment of the PDAC CAF signature in each murine pancreatitis fibroblast cluster. The PDAC CAF signature was obtained by comparing murine PDAC CAFs to pancreatitis fibroblasts from the snRNA-seq dataset. **(G)** Heatmap showing the fractional contribution of *in vivo* pancreatitis fibroblast subclusters to *in vivo* PDAC CAFs derived from averaged scANVI soft label probabilities. **(H)** Partition-based graph abstraction (PAGA) illustrates the inferred pseudotemporal trajectory among pancreatitis fibroblast subclusters and PDAC CAFs. Nodes (i.e., circles) represent cell subsets, and their size is scaled to the number of cells per cluster. Edges (i.e., line) represent the strength of connectivity (i.e., thickness). **(I)** Significantly upregulated and downregulated pathways identified by GSEA of PDAC CAFs compared to pancreatitis fibroblasts, as assessed by MAST from the snRNA-seq dataset. **(J)** Selected ligand–receptor interactions and their strength based on CellChat analysis between epithelial cells, fibroblasts, myeloids, and endothelial cells in pancreatitis and PDAC. Commun. Prob., communication probability. **(K)** Selected collagen pathway with different connections in pancreatitis and PDAC as assessed by CellChat analysis of the snRNA-seq datasets. **(L)** UMAP plot of epithelial cells in the combined snRNA-seq dataset from murine PDAC and pancreatitis. Different epithelial cell subsets are colour coded. **(M)** PAGA illustrating the inferred pseudotemporal trajectory among pancreatitis Ductal 1 and Ductal 2 cells and PDAC malignant cells. Nodes (i.e., circles) represent cell subsets, and their size is scaled to the number of cells per cluster. Edges (i.e., line) represent the strength of connectivity (i.e., thickness). **(N)** Significantly upregulated and downregulated pathways identified by GSEA of PDAC malignant cells compared to pancreatitis Ductal 1 or Ductal 2 cells, as assessed by MAST from the snRNA-seq dataset.

To further evaluate transcriptional changes between murine PDAC CAFs and pancreatitis fibroblasts, we looked for significantly altered pathways of the snRNA-seq dataset. In line with scRNA-seq of human tissues, this analysis showed an enrichment of myofibroblastic signatures (e.g., smooth muscle contraction, collagen formation) and myofibroblastic markers (e.g., *Thy1*, *Itga5*, *Ncam1*) in PDAC CAFs compared to pancreatitis fibroblasts (**Fig. 2I** and **1E**, **Extended Data Fig. 2F** and **Supplementary Table 2**)^4,5,7,27^. Inflammatory signatures (e.g. interferon response) were instead enriched in pancreatitis fibroblasts, and were highest in the *Pdgfra*^High^*Pi16*^High^ Fibro 2 and *Wt1*^High^*Dpp4*^High^ Fibro 4 subsets (**Fig. 2I** and **Extended Data Fig. 2H**). Consistent with our immunohistochemical analyses of tissues, snRNA-seq analysis also showed increased *Pdpn* and *Pdgfra* expression in pancreatitis fibroblasts and higher expression of *Acta2* (encoding αSMA) in PDAC CAFs (**Extended Data Fig. 2I and 1O;** **Fig. 1J** and **1N**). The expression of specific collagens varied between pancreatitis and PDAC fibroblasts, suggesting the presence of different ECM-producing profiles in pancreatitis and PDAC, as previously reported^12^ (**Extended Data Fig. 2I**). In keeping with this, CellChat analysis highlighted significant differences between ECM-associated ligands (e.g., collagens) involved in fibroblast-to-epithelial cell signalling in pancreatitis and PDAC (**Fig. 2J-K**).

Next, we aimed to determine whether fibroblasts from our mouse models of pancreatitis and PDAC recapitulate patterns of gene expression in these cells in humans. To do this, we leveraged the scRNA-seq dataset of human tissues and the snRNA-seq dataset of murine tissues and focused on curated protein-coding orthologs. First, we identified the 2,737 orthologs that had significantly higher expression (*P* adj < 0.05) in murine PDAC CAFs compared to pancreatitis fibroblasts (hereon, ‘murine PDAC CAF orthologs’). We also identified the 3,011 orthologs that had significantly higher expression in human PDAC CAFs compared to pancreatitis fibroblasts (hereon, ‘human PDAC CAF orthologs’). We found that 28% of human PDAC CAF orthologs (n=829/3,011) were also present in murine PDAC CAF orthologs (*P <* 1.9e-59, hypergeometric test; **Extended Data Fig. 2J** and **Supplementary Table 3**). Next, we performed cross-species comparison of pancreatitis fibroblast gene markers. We identified the 6,125 orthologs that had significantly higher expression in murine pancreatitis fibroblasts compared to PDAC CAFs (hereon, ‘murine pancreatitis fibroblast orthologs’) and the 6,461 orthologs that had significantly higher expression in human pancreatitis fibroblasts compared to PDAC CAFs (hereon, ‘human pancreatitis fibroblast orthologs’). We found that 54% (n=3,494/6,461) of human pancreatitis fibroblast orthologs overlapped with murine pancreatitis fibroblast orthologs (*P <* 3.4e-256; **Extended Data Fig. 2K** and **Supplementary Table 3**). Together, these data indicate that pancreatitis and PDAC mouse models significantly capture gene markers of human pancreatitis and PDAC fibroblasts.

Next, we looked to see if differential gene expression between epithelial cells in mouse pancreatitis and PDAC recapitulated differences in gene expression in these cells in humans. Analysis of the epithelial cell clusters from the snRNA-seq dataset of murine tissues identified two pancreatitis ductal cell populations, which we termed ‘Ductal 1’ and ‘Ductal 2’, and one PDAC malignant cell population (**Fig. 2L**). Comparison of the Ductal 1 and Ductal 2 clusters highlighted that the PDAC malignant cell signature (i.e., top 50 marker genes) was enriched in Ductal 1 compared to Ductal 2 cells, suggesting that Ductal 1 cells are transcriptomically closer to PDAC malignant cells (**Extended Data Fig. 2L** and **Supplementary Table 2**). This observation was further supported by pseudotemporal trajectory analysis that inferred a dominant trajectory connecting pancreatitis Ductal 1 cells to PDAC malignant cells (**Fig. 2M**). To further evaluate transcriptional differences between PDAC and pancreatitis epithelial cells, we compared the malignant cluster to the two pancreatitis ductal clusters, separately. PDAC malignant cells showed upregulation of proliferation-associated pathways (e.g., G2M checkpoint) compared to pancreatitis Ductal 2 cells (**Fig. 2N** and **Supplementary Table 2**). ECM-associated (e.g., ECM organisation) and inflammatory (e.g., NF-κB signalling) signatures were instead enriched in PDAC malignant compared to Ductal 1 cells (**Fig. 2N**). Thus, changes in murine PDAC malignant cells relative to pancreatitis ductal cells capture proliferation- and inflammation- associated signatures also observed in these cells in humans (**Fig. 1D**).

To further investigate the extent to which pancreatitis and PDAC epithelial cells in our mouse models recapitulate gene expression in these cells in humans, we used a similar approach to our cross-species comparison of gene expression in fibroblasts. First, we identified the 3,804 orthologs that had significantly higher expression in murine PDAC malignant compared to pancreatitis ductal cells (hereon, ‘murine PDAC malignant orthologs’), as well as the 1,751 orthologs that had significantly higher expression in human PDAC malignant compared to pancreatitis ductal cells (hereon, ‘human PDAC malignant orthologs’). We found that 43% (n=755/1,751) of human PDAC malignant orthologs were common to murine PDAC malignant orthologs (*P <* 7.1e-82; **Extended Data Fig. 2M** and **Supplementary Table 3**). Next, we identified the 5,378 orthologs that had significantly higher expression in murine pancreatitis ductal compared to PDAC malignant cells (hereon, ‘murine pancreatitis ductal orthologs’) and the 4,593 orthologs that had significantly higher expression in human pancreatitis ductal compared to PDAC malignant cells (hereon, ‘human pancreatitis ductal orthologs’). Comparison of these two datasets found that 47% (n=2,160/4,593) of human pancreatitis ductal orthologs were common to murine pancreatitis ductal orthologs (*P <* 5.0e-115; **Extended Data Fig. 2N** and **Supplementary Table 3**). These data indicate that pancreatitis and PDAC epithelial cells from mouse models significantly capture gene expression patterns of these cells in humans.

Together, these data reveal the presence of heterogeneous fibroblast and epithelial cell populations in pancreatitis, and highlight candidate precursors of PDAC CAFs and malignant cells, respectively. These results also support further comparative analyses of epithelial cells derived from our mouse models to investigate biology relevant to the respective human diseases.

### Pancreatitis-derived epithelial organoids capture *in vivo* gene expression patterns of pancreatitis ductal cells

PDAC epithelial organoid/PSC co-cultures have revolutionised our understanding of CAF heterogeneity, helping to identify iCAFs and myCAFs, and key malignant cell-produced mediators of their reprogramming^4,28^. We, thus, reasoned that establishing a similar model for pancreatitis could help identify distinct features of pancreatic fibroblasts and epithelial cells in inflammatory and malignant states. As a first step towards this, we generated pancreatitis epithelial organoids (hereon, ‘P organoids’) from the pancreata of C57BL/6J mice treated with caerulein for six weeks (**Fig. 3A**)^7,29^. We compared these P organoids to normal pancreatic organoids (hereon, ‘N organoids’) and PDAC organoids established from C57BL/6J mouse normal pancreata and KPC GEMM tumours, respectively (**Fig. 3A**). At day 1 post-isolation, as seen previously, N organoids displayed a budding ductal-like morphology and PDAC organoids formed large cystic cell clusters (**Fig. 3B**)^29^. In contrast, P organoids formed small cystic, relatively disorganised structures, suggesting they have a different origin to that of N organoids (**Fig. 3B**). After 2-3 passages in culture, the morphologies of N, P and PDAC organoids were similar, although P organoids grew slower than N and PDAC organoids (**Fig. 3C** and **Extended Data Fig. 3A-B**). In line with these results, RNA-sequencing (RNA-seq) analysis showed an upregulation of proliferation-associated signatures (e.g., G2M checkpoint) in N and PDAC compared to P organoids (**Fig. 3D-E** and **Extended Data Fig. 3C-D** and **Supplementary Table 4**). Additionally, as expected, N, P and PDAC organoids expressed genes markers of ductal epithelial cells (*Sox9*, *Pdx1*, *Krt19 –* encoding CK19; **Extended Data Fig. 3E** and **Supplementary Table 4**)^29^.

**Figure 3.**
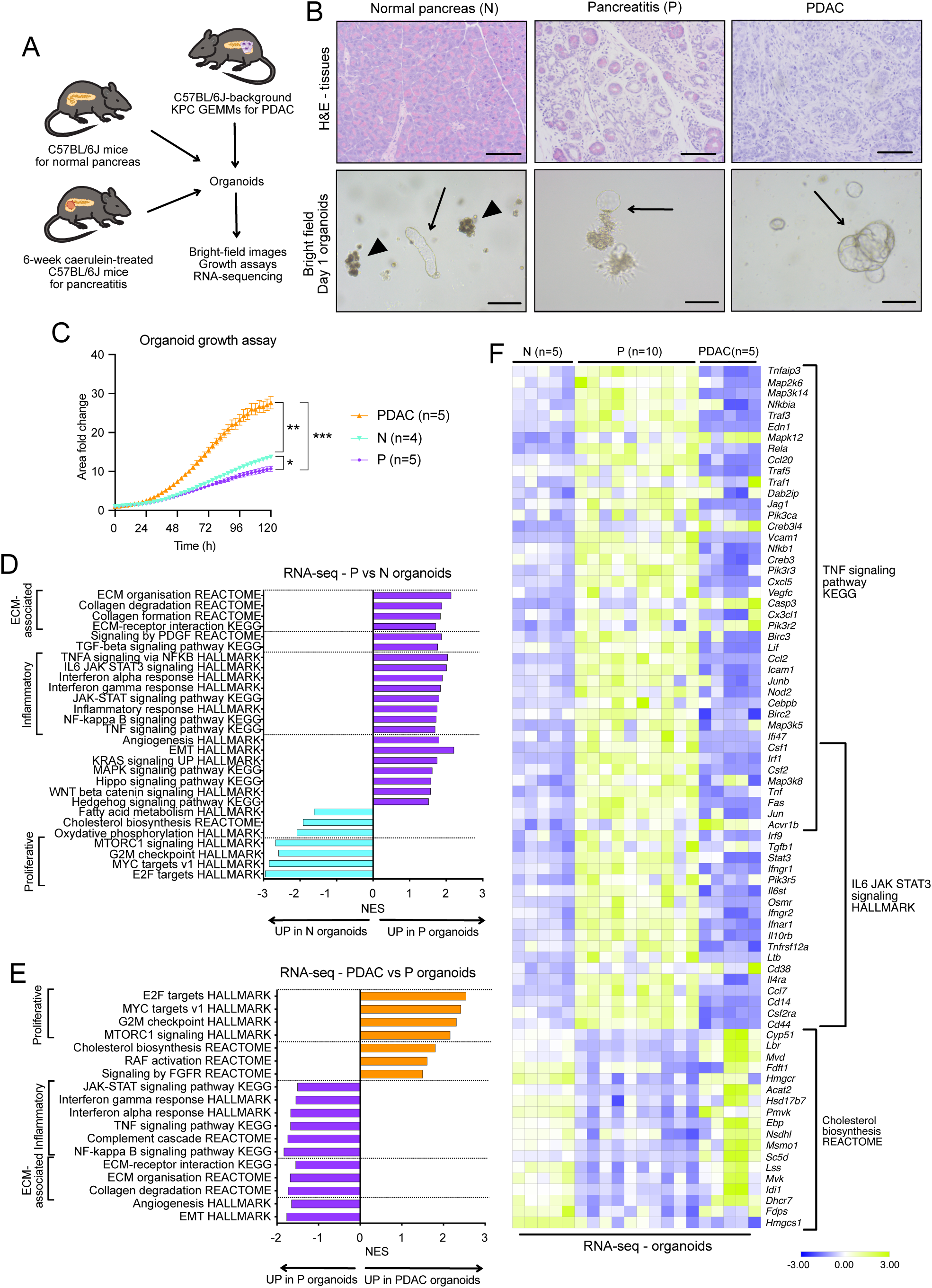
Pancreatitis-derived epithelial organoids capture *in vivo* gene expression patterns of pancreatitis ductal cells. **(A)** Schematic of models used for the generation of normal pancreas (i.e. from untreated mice), pancreatitis (i.e. from 6-week caerulein-treated mice) and PDAC (i.e. from KPC GEMMs) organoids, and techniques used for their analysis. **(B)** Top panels: Representative images of Haematoxylin & Eosin (H&E) stains of normal pancreas, pancreatitis and PDAC tissues; scale bars = 50 μm. Bottom panels: bright field images of epithelial organoids generated from normal pancreas (N), pancreatitis (P) and PDAC tissues at day 1 post-isolation; scale bars = 100 μm. Arrows indicate budding organoids. Triangles indicate clusters of acinar cells. **(C)** Proliferation curves of murine N, P and PDAC organoids in complete media. Results show mean ± SEM. * *P adj* < 0.05, ** *P adj* < 0.01, *** *P adj* < 0.001; Kruskal-Wallis test for the last time point. **(D)** Significantly upregulated and downregulated pathways identified by GSEA of P organoids (n=5, 2 different passages) compared to N organoids (n=5), as assessed by RNA-sequencing (RNA-seq). **(E)** Significantly upregulated and downregulated pathways identified by GSEA of PDAC organoids (n=5) compared to P organoids (n=5, 2 different passages), as assessed by RNA-seq. **(F)** Heatmap plot shows the normalised counts of selected genes and pathways upregulated or downregulated in P organoids compared to N and PDAC organoids are shown, as assessed by RNA-seq.

Further analysis of these organoid RNA-seq profiles revealed remarkable transcriptomic differences among N, P and PDAC organoids. P organoids upregulated ECM-associated (e.g., collagen degradation) and inflammatory (e.g., NF-κB and JAK-STAT signalling) pathways compared to N organoids (**Fig. 3D** and **3F**, **Extended Data Fig. 3F** and **Supplementary Table 4**). P organoids were enriched in ECM-associated (e.g., collagen degradation) and inflammatory (e.g., JAK-STAT signalling) pathways also relative to PDAC organoids (**Fig. 3E-F**, **Extended Data Fig. 3F** and **Supplementary Table 4**). In line with this, STAT3 activation (i.e., phospho-STAT3 levels) was higher in P compared to PDAC organoids (**Extended Data Fig. 3G**). Importantly, P organoids were also enriched for genes upregulated in pancreatitis Ductal 1 and Ductal 2 snRNA-seq profiles relative to PDAC malignant cells (**Extended Data Fig. 3H**). Furthermore, the PDAC malignant cell signature obtained from snRNA-seq profiles was enriched in PDAC organoids (**Extended Data Fig. 3I**).

These results demonstrate that P organoids retain features of pancreatitis ductal cells *in vivo* and suggest that these models could be leveraged to further investigate epithelial cell-fibroblast crosstalk in pancreatitis.

### The transcriptome of pancreatitis ductal cells is more influenced by fibroblast signalling compared to PDAC malignant cells

To investigate the crosstalk between epithelial cells and fibroblasts in PDAC and pancreatitis, we analysed five different culture conditions including: P organoids, PDAC organoids or PSCs alone (i.e., monocultures) to evaluate intrinsic patterns of gene expression in these cells when cultured in isolation; and P or PDAC organoids each co-cultured with PSCs to evaluate the impact of fibroblast signalling on pancreatitis and PDAC epithelial cell gene expression (**Fig. 4A-B** and **Extended Data Fig. 4A-B**). PSC numbers were significantly higher in P co-cultures compared to PDAC co-cultures or PSC monocultures, indicating that pancreatitis epithelial cells more effectively promote PSC expansion (**Fig. 4C**). In keeping with immunohistochemical and snRNA-seq analyses of murine tissues, P co-cultures also displayed higher fibroblast-to-epithelial cell ratios compared to PDAC co-cultures (**Fig. 4D** and **Extended Data Fig. 4C-D** and **Supplementary Table 2**). These results suggest that co-culturing P and PDAC organoids with PSCs can recapitulate pancreatitis and PDAC epithelial cell-fibroblast composition found *in vivo*.

**Figure 4.**
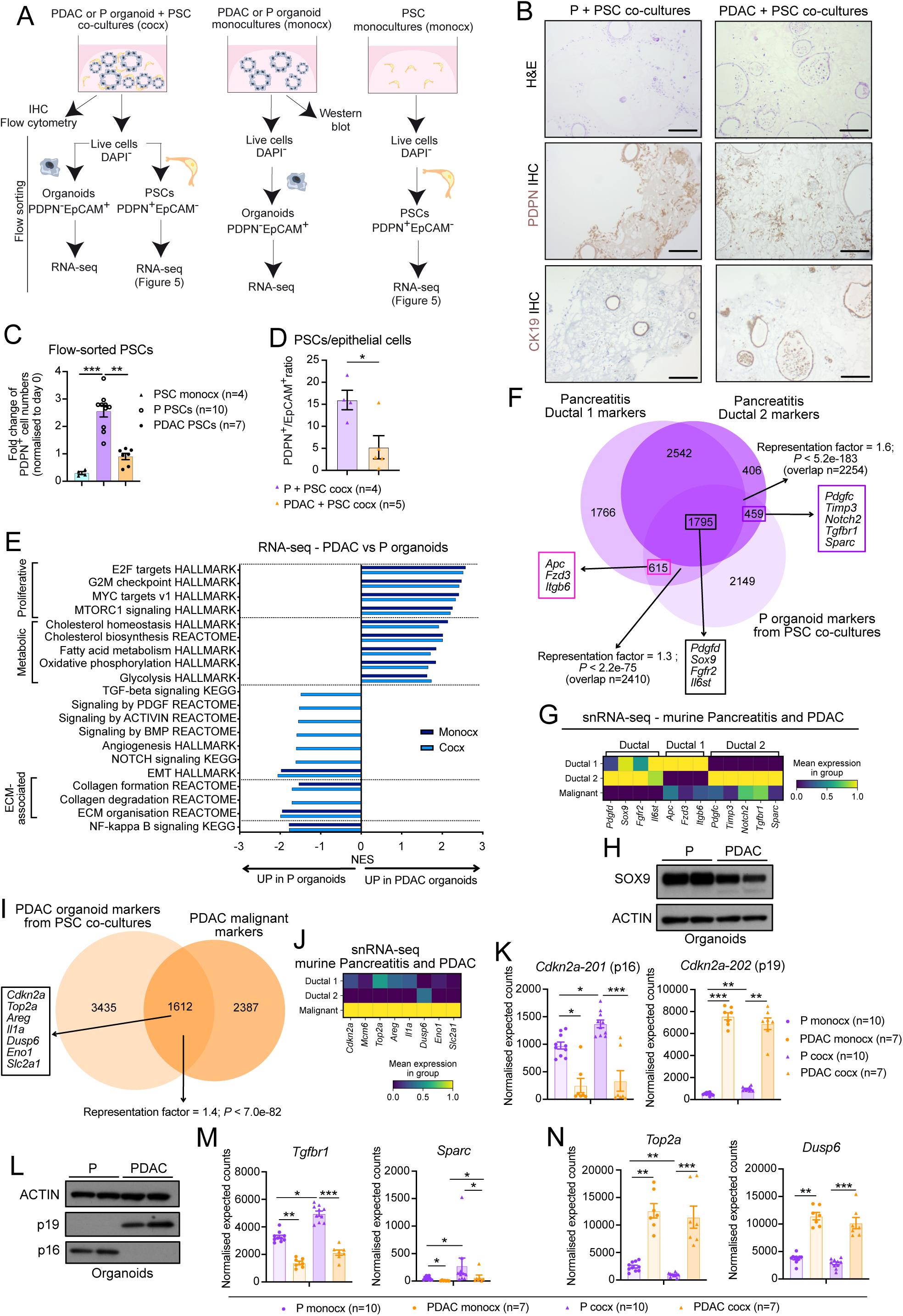
The transcriptome of pancreatitis ductal cells is more influenced by fibroblast signalling compared to PDAC malignant cells. **(A)** Schematic of flow sorting strategy of organoid and pancreatic stellate cell (PSC) monocultures (monocx) and P or PDAC organoid/PSC co-cultures (cocx) in reduced media (i.e. 5% FBS/DMEM) prior to RNA-seq. **(B)** Representative images of H&E, podoplanin (PDPN) and CK19 stains of P or PDAC organoid/PSC cocx after 7 days in culture. Scale bars = 100 μm. **(C)** Fold change in numbers of PSCs (PDPN^+^EpCAM^-^) flow-sorted from monocultures (monocx) or from P or PDAC cocx after 3.5 days in culture relative to numbers of PSCs plated at day 0. Results show mean ± SEM. ** *P adj* < 0.01; *** *P adj* < 0.001, Kruskal-Wallis test. **(D)** Fibroblast (PDPN^+^EpCAM^-^)/epithelial cell (EpCAM^+^PDPN^-^) ratio as assessed by flow cytometry of P or PDAC organoid/PSC cocx after 3.5 days in culture. Results show mean ± SEM. * *P* < 0.05, Mann-Whitney test. **(E)** Significantly upregulated (i.e., normalised enrichment score, NES > 1.50 and FDR < 0.25) and downregulated (i.e., NES < -1.50 and FDR < 0.25, apart for TGF-β-signaling KEGG with NES = -1.496) pathways identified by GSEA of PDAC organoids in monocx or cocx compared to P organoids in monoculture or co-culture, respectively, as assessed by RNA-seq. **(F)** Venn diagrams of curated protein-coding genes significantly (*P* adj < 0.05) upregulated in P organoids in PSC co-culture compared to PDAC organoids in PSC co-culture (hereon, ‘P organoid markers from PSC co-cultures) and murine pancreatitis Ductal 1 or Ductal 2 cells compared to PDAC malignant cells *in vivo* (hereon, ‘pancreatitis Ductal 1 markers’ and ‘pancreatitis Ductal 2 markers’, respectively), as assessed by RNA-seq and snRNA-seq, respectively. Significance of the overlap between datasets was defined by hypergeometric test (with denominator n=18,000). Selected genes common to two or more datasets are indicated. **(G)** Heatmap of scaled expression of selected genes differentially upregulated in murine pancreatitis epithelial clusters. Data are scaled such that the cluster with the lowest average expression = 0 and the highest = 1 for each gene. **(H)** Western blot analysis of SOX9 in P and PDAC organoids cultured in reduced media for 3 days. ACTIN, loading control. **(I)** Venn diagrams of curated protein-coding genes significantly (*P* adj < 0.05) upregulated in murine PDAC malignant cells compared to pancreatitis ductal cells *in vivo* (hereon, ‘PDAC malignant markers’) and murine PDAC organoids in PSC co-culture compared murine P organoids in PSC co-culture (hereon, ‘PDAC organoid markers from PSC co-cultures’), as assessed by snRNA-seq and RNA-seq, respectively. Significance of the overlap between datasets was defined by hypergeometric test (with denominator n=18,000). Selected genes common to two or three datasets are indicated. **(J)** Heatmap of scaled expression of genes differentially upregulated in murine PDAC epithelial clusters. Data are scaled such that the cluster with the lowest average expression = 0 and the highest = 1 for each gene. **(K)** RNA-seq expression of *Cdkn2a-201* (encoding p16) and *Cdkn2a-202* (encoding p19) in P or PDAC monocx and cocx with PSCs. Results show mean ± SEM. * *P adj* < 0.05; ** *P adj* < 0.01; *** *P adj* < 0.001, Kruskal-Wallis test. **(L)** Western blot analysis of p19 and p16 in P and PDAC organoids cultured in reduced media for 3 days. ACTIN, loading control. **(M)** RNA-seq expression of *Tgfbr1* and *Sparc* in P or PDAC organoid monocx and P or PDAC cocx with PSCs. Results show mean ± SEM. * *P adj* < 0.05; ** *P adj* < 0.01; *** *P adj* < 0.001, Kruskal-Wallis test. **(N)** RNA-seq expression of *Top2a* and *Dusp6* in P or PDAC monocx and cocx with PSCs. Results show mean ± SEM. ** *P adj* < 0.01; *** *P adj* < 0.001, Kruskal-Wallis test.

Consistent with what we observed by transcriptomic analyses of epithelial cells from human and murine tissues, proliferation-associated signatures (e.g., G2M checkpoint) were enriched in PDAC compared to P organoids (**Fig. 4E, 1D and 2N**; **Extended Data Fig. 4E-F**; **Supplementary Table 5**). Indeed, these signatures were enriched in PDAC relative to P organoids independent of culture condition (**Fig. 4E**). In contrast, pathways upregulated in P compared to PDAC organoids appeared to be largely dependent on the presence of PSCs in co-cultures (**Fig. 4E**). These pathways included PDGF, transforming growth factor beta (TGF-β) and NOTCH signalling, which are known to mediate key epithelial-stromal crosstalk^2^. Additional analyses revealed that while both P and PDAC organoids upregulated certain inflammatory and ECM-associated pathways in co-cultures compared to their respective monocultures, these enrichments were highest in P organoid co-cultures (**Extended Data Fig. 4G-H** and **Supplementary Table 5**). Together, these findings suggest that while both P and PDAC organoids respond to PSC-derived signals, distinct epithelial–stromal cross-talks operate in pancreatitis and PDAC. Finally, the Ductal 1, but not Ductal 2, signature from murine pancreatitis snRNA-seq profiles was enriched in P organoids from co-cultures relative to P monocultures, suggesting that the pancreatitis Ductal 1 state found *in vivo* may be partially dependent on stromal signalling (**Extended Data Fig. 4I**).

To further determine whether the crosstalk between PSCs and P or PDAC organoids in co-cultures promoted epithelial cell gene expression patterns similar to those observed in murine pancreatitis and PDAC tissues, we leveraged an analogous strategy to what we used for murine and human tissue comparisons. First, we identified the genes that had significantly higher expression in P compared to PDAC organoid co-cultures (hereon, ‘P organoid markers from PSC co-cultures’), as well as the two murine gene sets that were expressed at significantly higher levels in Ductal 1 and Ductal 2 snRNA-seq profiles compared to PDAC malignant cells (hereon, ‘pancreatitis Ductal 1 markers’ and ‘pancreatitis Ductal 2 markers’, respectively). We found that P organoid markers from PSC co-cultures significantly overlapped with both pancreatitis Ductal 1 and Ductal 2 markers (*P <* 2.2e-75 and *P <* 5.2e-183, respectively; **Fig. 4F** and **Supplementary Table 3**). This analysis also enabled the identification of genes that were upregulated in both Ductal 1 and Ductal 2 cells, only in Ductal 1 cells or only in Ductal 2 cells compared to PDAC malignant cells (**Fig. 4F-G**). In line with the pathways enriched in P compared to PDAC organoid co-cultures, these genes included ligands and receptors involved in PDGF (e.g., *Pdgfd*, *Pdgfc*), TGF-β (e.g., *Tgfbr1*, *Sparc*) and NOTCH (e.g., *Notch2*) signalling (**Fig. 4E-G**).

Next, we aimed to determine whether P organoids from PSC co-cultures also recapitulate gene patterns of ductal cells found in human pancreatitis tissues. We identified the orthologs of the P organoid markers from PSC co-cultures (hereon, ‘P organoid orthologs from PSC co-cultures’) and compared them with the human pancreatitis ductal orthologs previously identified (**Extended Data Fig. 4J and 2N**). This analysis showed that 39% (n=1,777/4,593) of human pancreatitis ductal orthologs were common to P organoid orthologs from PSC co-cultures (*P <* 5.1e-74; **Extended Data Fig. 4J**). Furthermore, 55% (n=977/1,777) of these markers were also common to the murine pancreatitis ductal orthologs previously identified (e.g., *SOX9*, *PDGFD*, *FGFR2*, *IL6ST*; **Extended Data Fig. 4J and 2N**). Among these genes, SOX9 has been previously described as a marker of ADM and to be expressed in PDAC^30,31^. We validated that the SOX9 protein was upregulated in P compared to PDAC organoids (**Fig. 4H**). Notably, this cross-species, cross-model comparison also enabled the identification of 800 human pancreatitis ductal orthologs that were common to P organoid orthologs from PSC co-cultures but not to murine pancreatitis ductal orthologs (e.g., *STAT3*, *NFKB1*, *MMP24*; **Extended Data Fig. 4J**). These data suggest that analysis of P organoid PSC co-cultures can complement analysis of pancreatitis mouse models to dissect gene patterns of the human disease.

We then aimed to determine whether PDAC organoids from PSC co-cultures recapitulate gene expression profiles of malignant cells from PDAC tissues. First, we identified the genes that had significantly higher expression in PDAC compared to P organoid co-cultures (hereon, ‘PDAC organoid markers from PSC co-cultures’) and the genes that had significantly higher expression in murine PDAC malignant snRNA-seq profiles compared to pancreatitis ductal cells (hereon, ‘murine PDAC malignant markers’). We found a significant overlap between PDAC organoid markers from PSC co-cultures and murine PDAC malignant markers (*P <* 7.0e-82; **Fig. 4I-J** and **Supplementary Table 3**). Next, we identified the orthologs of PDAC organoid markers from PSC co-cultures (hereon, ‘PDAC organoid orthologs from PSC co-cultures’) and compared them with the human PDAC malignant orthologs previously identified (**Extended Data Fig. 2M**). We found that 46% (n=798/1,751) of human PDAC malignant orthologs were common to both datasets (*P <* 5.7e-57; **Extended Data Fig. 4K**). Of these, 47% (n=372/798) were also upregulated in the murine PDAC malignant orthologs previously identified (**Extended Data Fig. 4K** and **2M**). In keeping with the pathways enriched in PDAC compared to P organoids, among the orthologs shared across all species and models were proliferative (e.g., *TOP2A*) and metabolic (e.g., *SLC2A1*) markers (**Extended Data Fig. 4K and Fig. 4E**). These shared orthologs also encoded markers previously found to be abundant in PDAC, including *GCNT3* and *MUC5AC*^29^. Intriguingly, the expression of *CDKN2A*, which encodes p16 and p14 proteins through alternative reading frames in humans and is frequently lost in PDAC^32^, was also upregulated in PDAC malignant compared to pancreatitis ductal cells *in vitro* and *in vivo* (**Fig. 4I-J** and **Extended Data Fig. 4K**). Differential expression analysis of the separate transcripts revealed that *Cdkn2a-202* (encoding p19 - the murine ortholog of p14), but not *Cdkn2a-201* (encoding p16), was upregulated in PDAC compared to P organoids (**Fig. 4K** and **Supplementary Table 5**). Protein analysis confirmed upregulation of p19, but not p16, in PDAC organoids (**Fig. 4L**). Our (KPC) PDAC organoids are driven by *Kras*^G12D^ and *Trp53*^R172H^ mutations and have undergone loss of heterozygosity (LOH) of the *Trp53* wild-type allele^33^. To determine whether the presence of mutant KRAS and/or mutant p53 led to p19 upregulation in PDAC organoids, we compared p19 protein abundance in N and PDAC organoids, as well as previously characterised KC (*Kras*^LSL-G12D/+^; *Pdx1*-Cre) GEMM-derived pancreatic intraepithelial neoplasia organoids (hereon, ‘KC organoids’) and KPC PDAC organoids that have not undergone p53 LOH (hereon, ‘KPC organoids’)^33^. This analysis showed that while p19 levels increased from N to KC organoids, they were comparable between KC and KPC organoids, and were highest in PDAC organoids with p53 LOH (**Extended Data Fig. 4L**). These results suggest that both KRAS^G12D^ and p53 LOH lead to p19 upregulation in PDAC malignant cells compared to pancreatitis ductal cells.

PSCs appeared to impact the gene expression in P organoids from co-cultures far more than in PDAC organoids (**Fig. 4E**). In line with this, while the expression of only 1,042 genes differed significantly between PDAC organoid mono- and co-cultures, more than three times as many genes (n=3,389) were significantly altered in expression between P mono- and co-cultures (**Supplementary Table 5**). We then aimed to determine what proportion of P and PDAC organoid markers from PSC co-cultures are regulated by PSC-derived cues. Simply comparing co-cultures does not reveal which features are intrinsic to organoids and which are influenced by PSCs. To address this, we compared the P organoid markers from PSC co-cultures with genes significantly upregulated in P organoids when co-cultured versus in monoculture. This analysis identified 20% (n=1,016/5,018) of P organoid markers from PSC co-cultures that were also upregulated in P organoids by PSC-derived cues (e.g., *Tgfbr1*, *Sparc*; **Fig. 4M**, **Extended Data Fig. 4M** and **Supplementary Table 5**). Next, we compared the PDAC organoid markers from PSC co-cultures with genes significantly upregulated in PDAC organoids when co-cultured versus in monoculture. This analysis identified only 3% (n=138/5,047) of PDAC organoid markers from PSC co-cultures that were also upregulated in PDAC organoids by PSC-derived cues (**Extended Data Fig. 4N** and **Supplementary Table 5**). Indeed, the expression of most PDAC organoid markers from PSC co-cultures (e.g., *Top2a*, *Dusp6*) was not significantly different between mono- and co-cultures (**Fig. 4N**).

Together, these data indicate that P and PDAC organoid/PSC co-cultures recapitulate *in vivo* gene expression patterns of murine and human epithelial cells in pancreatitis and PDAC. These results also suggest that the transcriptome of pancreatitis ductal cells is more influenced by fibroblast signalling compared to PDAC malignant cells.

### Organoid/PSC co-cultures only partially recapitulate the *in vivo* transcriptomic differences of pancreatitis and PDAC fibroblasts

We next aimed to understand how pancreatitis and PDAC epithelial cells differentially reprogram fibroblasts. To do this, we analysed the transcriptome of PSCs from monocultures and from co-cultures with P or PDAC organoids (**Fig. 4A**). We found that PSCs were activated in both P and PDAC co-cultures compared to PSC monocultures, as indicated by a reduction in quiescent fibroblast markers (*Plin4*, *Kitl*) and an increase in inflammatory (*Il6*, *Cxcl1*) and myofibroblastic (*Tgfb1*, *Col1a1*) markers (**Fig. 5A-B** and **Extended Data Fig. 5A**)^4,13,28,34^. In line with these results, inflammatory (e.g., NF-κB and JAK/STAT signalling) and myofibroblastic (e.g., collagen formation) signatures were enriched in both PSC co-cultures compared to PSC monocultures (**Extended Data Fig. 5B** and **Supplementary Table 6**)^28,35^. Despite these similarities, inflammatory markers (*Il1a*, *Il6*, *Cxcl1*) and pathways (e.g., NF-κB signalling, interferon response) were higher in PSCs from PDAC co-cultures (**Fig. 5A and 5C-D**; **Supplementary Table 6**)^28^. To identify candidate mediators of this increased fibro-inflammatory phenotype, we evaluated the expression of known epithelial cell-derived ligands that induce inflammatory fibroblast and myofibroblast phenotypes in PDAC. The expression of *Tgfb1*, which mediates myCAF formation, was comparable in P and PDAC organoids (**Fig. 5E**)^28^. On the contrary, *Il1a* and *Il1b* (encoding interleukin 1 alpha and beta, respectively), which mediate iCAF formation, had significantly higher expression in PDAC organoids from co-cultures (**Fig. 5E** and **Extended Data Fig. 5C**)^28^. In keeping with these results, NicheNet analysis inferred *Il1a* and *Il1b* as candidate mediators of the PDAC organoid-to-PSC crosstalk (**Extended Data Fig. 5D-E**). Thus, increased IL-1 signalling may mediate the more pronounced inflammatory state of PSCs from PDAC co-cultures compared to P co-cultures.

**Figure 5.**
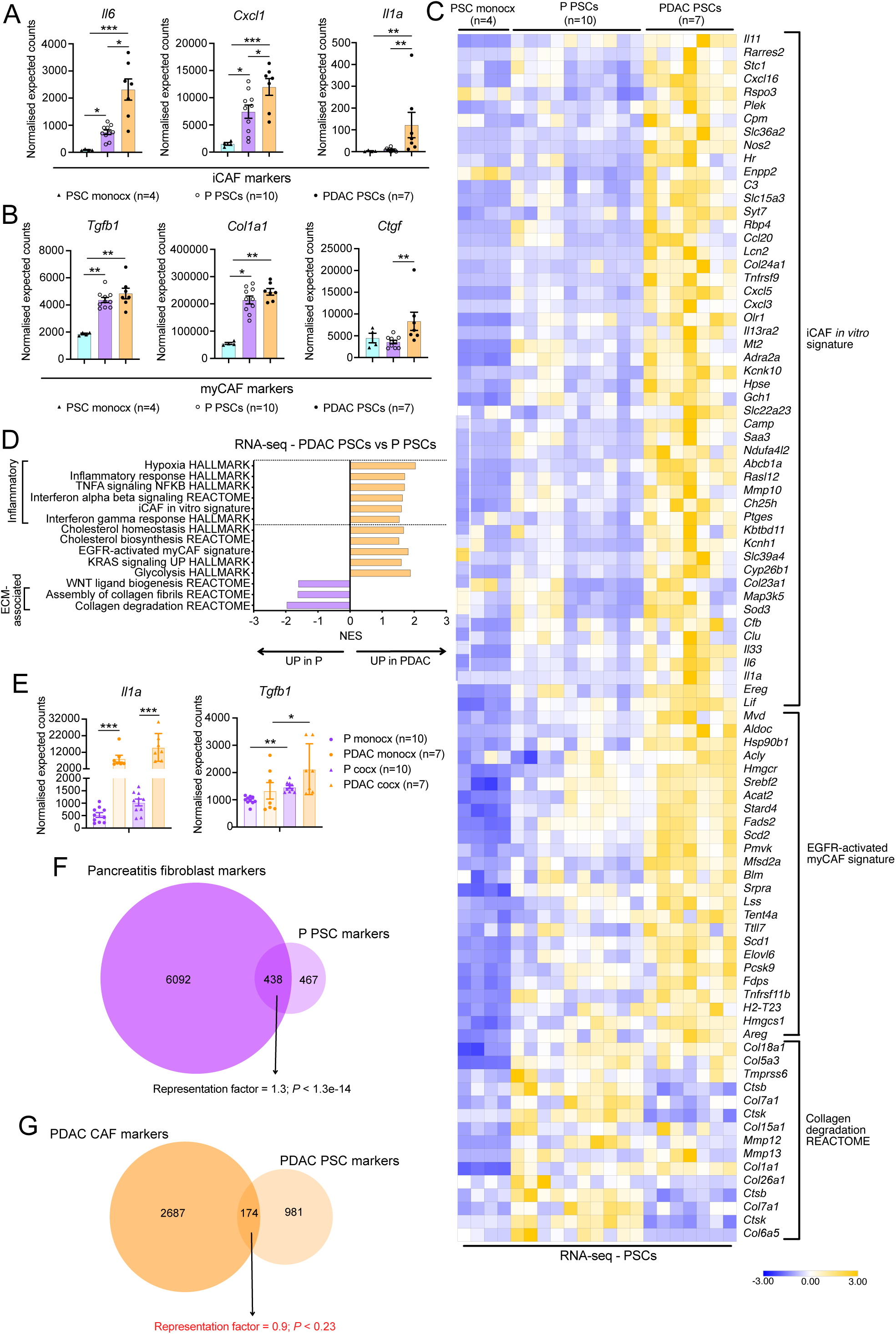
Organoid/PSC co-cultures only partially recapitulate the *in vivo* transcriptomic differences of pancreatitis and PDAC fibroblasts. (A) RNA-seq expression of iCAF markers (*Il1a*, *Il6*, *Cxcl1*) in PSC monocultures (n=2, 2 different passages) and co-cultures with P (n=10) or PDAC (n=7) organoids. Results show mean ± SEM. * *P* < *adj* 0.05; ** *P adj* < 0.01; *** *P adj* < 0.001, Kruskal-Wallis test. **(B)** RNA-seq expression of myCAF markers (*Tgfb1*, *Col1a1*, *Ctgf*) in PSC monocultures and co-cultures with P or PDAC organoids. Results show mean ± SEM. ** *P adj* < 0.01, Kruskal-Wallis test. **(C)** Heatmap plot shows the normalised counts from RNA-seq of murine PSC monocultures, PSCs cultured with P organoids and PSCs cultured with PDAC organoids. Selected genes and pathways upregulated or downregulated across conditions are shown. **(D)** Significantly upregulated and downregulated pathways identified by GSEA of PSCs cultured with PDAC organoids compared to PSCs cultured with P organoids, as assessed by RNA-seq. **(E)** RNA-seq expression of *Il1a* and *Tgfb1* in P or PDAC organoid monocultures and co-cultures with PSCs. Results show mean ± SEM. * *P adj* < 0.05, ** *P adj* < 0.01; *** *P adj* < 0.001, Kruskal-Wallis test. **(F)** Venn diagrams of curated protein-coding genes significantly (*P* adj < 0.05) upregulated in murine PSCs cultured with P organoids compared to murine PSCs cultured with PDAC organoids (hereon, ‘P PSC markers’), and murine pancreatitis fibroblasts compared to PDAC CAFs *in vivo* (hereon, ‘pancreatitis fibroblast markers’), as assessed by RNA-seq and snRNA-seq, respectively. Significance of the overlap between datasets was defined by hypergeometric test (with denominator = 18,000). **(G)** Venn diagrams of curated protein-coding genes significantly (*P* adj < 0.05) upregulated in murine PSCs cultured with PDAC organoids compared to murine PSCs cultured with P organoids (hereon, ‘PDAC PSC markers’), and murine PDAC CAFs compared to murine pancreatitis fibroblasts *in vivo* (hereon, ‘PDAC CAF markers’), as assessed by RNA-seq and snRNA-seq, respectively. Significance of the overlap between datasets was defined by hypergeometric test (with denominator = 18,000).

Similar to our analysis of epithelial cells, we next looked to see if P or PDAC organoids impacted gene expression in PSCs in a manner that recapitulated gene expression profiles of fibroblasts in pancreatitis and PDAC tissues. First, we identified the genes significantly upregulated in PSCs from P co-cultures compared to PSCs from PDAC co-cultures (hereon, ‘P PSC markers’) and the genes significantly upregulated in murine pancreatitis fibroblast snRNA-seq profiles compared to PDAC CAFs (hereon, ‘pancreatitis fibroblast markers’). We found a significant overlap between these two datasets (*P <* 1.3e-14; **Fig. 5F** and **Supplementary Table 3**), as well as between the orthologs of the P PSC markers (hereon, ‘P PSC orthologs’) and the human pancreatitis fibroblast orthologs previously identified (*P <* 1.2e-10; **Extended Data Fig. 5F and 2K**; **Supplementary Table 3**). Next, we aimed to determine whether PSCs cultured with PDAC organoids similarly captured *in vivo* gene markers of PDAC CAFs. To do this, we identified the genes significantly upregulated in PSCs from PDAC co-cultures compared to PSCs from P co-cultures (hereon, ‘PDAC PSC markers’), as well as the genes significantly upregulated in murine PDAC CAF snRNA-seq profiles compared to pancreatitis fibroblasts (hereon, ‘PDAC CAF markers’). We found that these two datasets did not significantly overlap (*P* < 0.23; **Fig. 5G** and **Supplementary Table 3**). Orthologs of the PDAC PSC markers (hereon, ‘PDAC PSC orthologs’) also did not significantly overlap with the human PDAC CAF orthologs previously identified (*P <* 0.21; **Extended Data Fig. 5G and 2J**; **Supplementary Table 3**). This limited overlap in gene expression profiles between PDAC PSCs and *in vivo* CAFs is in line with the observation that PSCs from PDAC co-cultures were not enriched in pathways (e.g., myofibroblastic signatures) that were instead enriched in PDAC CAFs from murine and human tissues when compared to their respective pancreatitis fibroblasts (**Fig. 5D**, **1E** and **2I**).

Thus, P and PDAC organoid/PSC co-cultures only partially recapitulate the *in vivo* gene expression differences observed in fibroblasts from pancreatitis and PDAC tissues. Incorporating additional stromal cell types may be necessary to better capture the phenotypic differences seen *in vivo*.

### Cross-model analysis pinpoints epithelial cell markers of pancreatitis and PDAC

Previous work showed that PSCs only give rise to a subset of PDAC CAFs, and that other CAF precursors include pancreatic fibroblasts and mesothelial cells^6,8,14,36–38^. We reasoned that pancreatitis fibroblasts may similarly have different cells of origin. Indeed, our snRNA-seq dataset of murine pancreatitis tissues revealed the presence of four fibroblast subsets, and two of these subsets (Fibro 1 and Fibro 4) expressed mesothelial cell markers (e.g., *Wt1*; **Fig. 2E** and **Extended Data Fig. 2E**). These considerations prompted us to optimise our organoid co-culture models to better mimic the heterogeneity of pancreatitis and PDAC fibroblasts *in vivo*.

First, we established and characterised fibroblast and mesothelial cell lines from normal pancreata of C57BL/6J mice (**Extended Data Fig. 6A**)^4^. In 2D cultures, fibroblasts displayed an elongated morphology similar to PSCs while mesothelial cells were isolated by enriching for cells with cobblestone-like morphology, which is characteristic of mesothelial cells in 2D culture (**Extended Data Fig. 6A-B**)^39,40^. Both fibroblast and mesothelial cell cultures lacked the expression of markers of immune (*Ptprc*), endothelial (*Pecam1*), pericyte (*Rgs5*) and epithelial (*Epcam*) cells (**Extended Data Fig. 6C-D**). Furthermore, fibroblast and mesothelial cell cultures expressed significantly greater levels of fibroblastic (*Acta2*, *Col1a1*, *Pdgfra* and *Pi16*) and mesothelial (*Msln*, *Wt1*, *Krt19*) markers, respectively (**Extended Data Fig. 6E**)^38^. Mesothelial cells also expressed higher cell-surface levels of the mesothelial cell protein CD200 relative to PSCs (**Extended Data Fig. 6F**). Finally, since a feature of mesothelial cells is their ability to present major histocompatibility complex type II (MHCII) upon stimulation, we assessed MHCII cell-surface levels of interferon-γ (IFNγ)-treated PSCs, fibroblasts and mesothelial cells^41^. Upon IFNγ stimulation, fibroblasts and mesothelial cells presented significantly higher cell-surface levels of MHCII relative to PSCs (**Extended Data Fig. 6G**). Together these results show that fibroblast lines represent mixed populations of fibroblasts, mesothelial cells and, potentially, PSCs.

We then aimed to evaluate how the transcriptomes of P and PDAC organoids was differently shaped in co-culture with PSCs, fibroblasts and mesothelial cells (hereon, ‘multi-stroma’). To do this, we performed scRNA-seq analysis of multi-stromal organoid models comprised of green fluorescent protein-positive (GFP^+^) PSCs, TdTomato^+^ fibroblasts, TdTomato^+^ mesothelial cells and P or PDAC organoids (**Fig. 6A-C** and **Extended Data Fig. 6H-M**). In keeping with our snRNA-seq analysis of murine tissues, we identified two P organoid clusters (hereon, ‘P organoid 1’ and ‘P organoid 2’) and one PDAC organoid cluster (**Fig. 6C** and **2C**). Also in line with our snRNA-seq analysis, the PDAC malignant signature found in murine tissues was upregulated in PDAC compared to P organoids, as well as in P organoid 1 compared to P organoid 2 cells, from multi-stroma co-cultures (**Extended Data Fig. 6N-O**). Furthermore, markers specific to pancreatitis Ductal 1 cells from murine tissues were enriched in P organoid 1 compared to P organoid 2 cells from multi-stroma co-cultures (**Extended Data Fig. 6O**). Thus, we evaluated whether pathways enriched in PDAC malignant cells relative to pancreatitis Ductal 1 and Ductal 2 cells from murine tissues were also enriched in PDAC organoids relative to P organoid 1 and P organoid 2 cells from multi-stroma co-cultures. We found that proliferation-associated pathways (e.g., G2M checkpoint) were upregulated in PDAC organoid compared to P organoid 2 cells (**Fig. 6D** and **Supplementary Table 7**). ECM-associated (e.g., ECM organisation) and inflammatory (e.g., NF-κB signalling) signatures were instead enriched in PDAC organoid compared to P organoid 1 cells (**Fig. 6D**). Thus, changes in PDAC organoids compared to P organoids in multi-stromal co-cultures mirror changes observed in PDAC malignant cells relative to pancreatitis ductal cells from murine and human tissues (**Fig. 1D** and **2N**).

**Figure 6.**
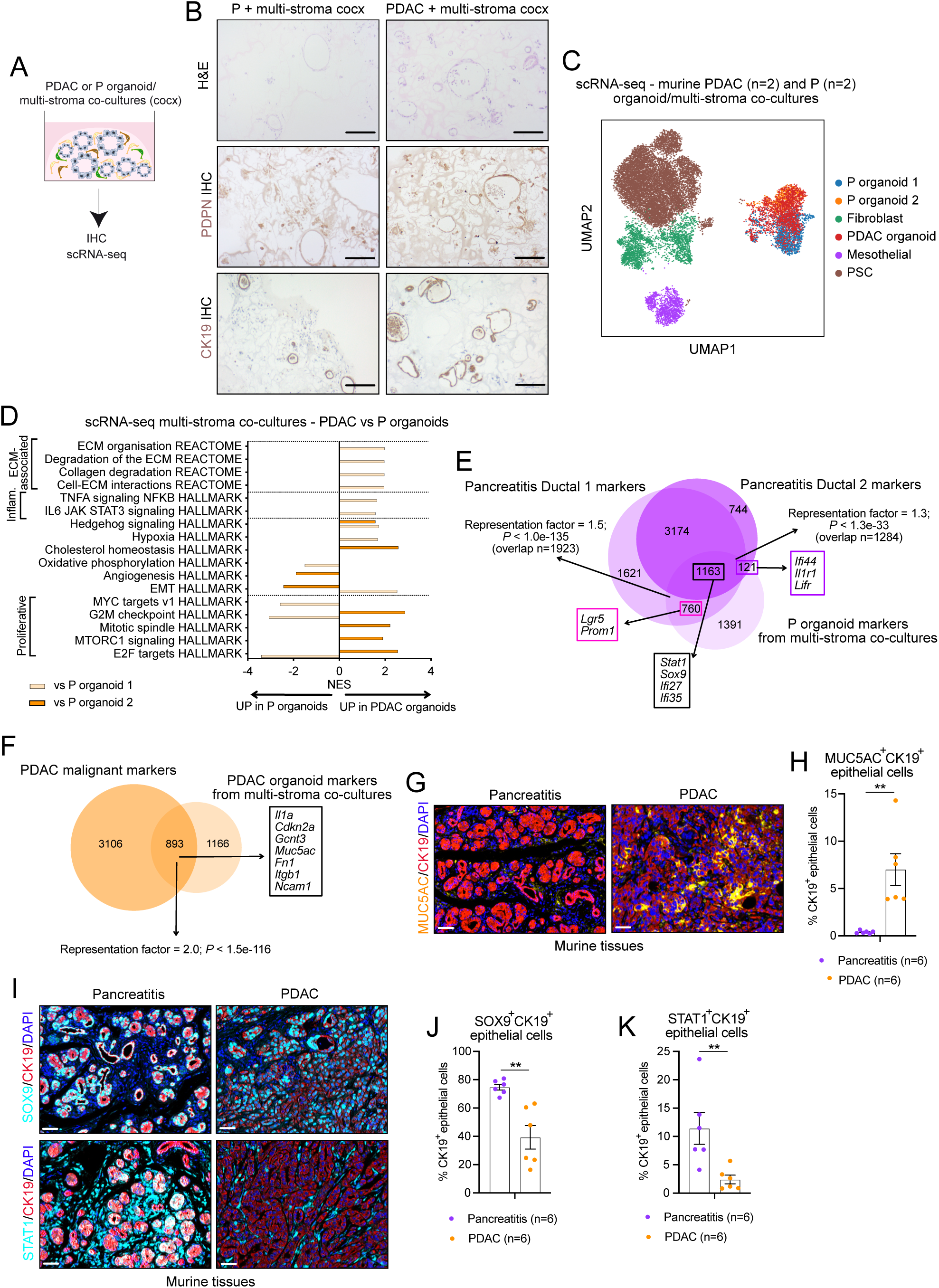
Cross-model analysis pinpoints epithelial cell markers of pancreatitis and PDAC. **(A)** Schematic of analyses of P or PDAC organoid/multi-stroma co-cultures in reduced media (i.e. 5% FBS/DMEM). **(B)** Representative images of H&E, PDPN and CK19 stains of P or PDAC organoid/multi-stroma co-cultures after 7 days in culture. Scale bars = 100 μm. **(C)** UMAP plot of cell subsets in the combined scRNA-seq dataset of PDAC (n=2) and P (n=2) organoid/multi-stroma co-cultures after 3.5 days in culture. Different cell subsets are colour coded. **(D)** Significantly upregulated and downregulated pathways identified by GSEA of PDAC organoids compared to P organoids 1 or P organoids 2, as assessed by MAST from the scRNA-seq dataset of organoid/multi-stroma co-cultures. **(E)** Venn diagrams of curated protein-coding genes significantly (*P* adj < 0.05) upregulated in P organoids in multi-stroma co-culture compared to PDAC organoids in multi-stroma co-culture (hereon, ‘P organoid markers from multi-stroma co-cultures’), as assessed by scRNA-seq, and pancreatitis Ductal 1 or Ductal 2 markers. Significance of the overlap between datasets was defined by hypergeometric test (with denominator n=18,000). Selected genes common to all datasets are indicated. **(F)** Venn diagrams of curated protein-coding genes significantly upregulated in PDAC organoids in multi-stroma co-culture compared to P organoids in multi-stroma co-culture (hereon, ‘PDAC organoid markers from multi-stroma co-cultures’), as assessed by scRNA-seq, and PDAC malignant markers. Significance of the overlap between datasets was defined by hypergeometric test (with denominator n=18,000). Selected genes common to all datasets are indicated. **(G)** Representative multiplex immunofluorescence (IF) images of MUC5AC (yellow), CK19 (red) and DAPI (nuclear stain, blue) in murine pancreatitis and PDAC tissues. Scale bars = 50 μm. **(H)** Quantification of MUC5AC^+^CK19^+^DAPI^+^αSMA^-^PDGFRα^-^ cells relative to DAPI^+^CK19^+^αSMA^-^PDGFRα^-^ epithelial cells in murine pancreatitis and PDAC tissues, as assessed by multiplex IF analysis. Results show mean ± SEM. ** *P* < 0.01, Mann-Whitney test. **(I)** Representative multiplex IF images of SOX9 (cyan), CK19 (red) and DAPI (nuclear stain, blue) (top panels), and STAT1 (cyan), CK19 (red) and DAPI (blue) (bottom panels) in murine pancreatitis and PDAC tissues. Scale bars = 50 μm. **(J-K)** Quantification of SOX9^+^CK19^+^DAPI^+^ **(J)** and STAT1^+^CK19^+^DAPI^+^ **(K)** αSMA^-^PDGFRα^-^ cells relative to DAPI^+^CK19^+^αSMA^-^PDGFRα^-^ epithelial cells in murine pancreatitis and PDAC tissues, as assessed by multiplex IF analysis. Results show mean ± SEM. ** *P* < 0.01, Mann-Whitney test.

To further investigate the degree of overlap between gene markers of P and PDAC organoids in multi-stroma co-cultures and gene markers of pancreatitis and PDAC epithelial cells in tissues, we leveraged similar strategies used for cross-model and cross-species comparisons with organoid/PSC co-cultures. For murine comparisons, we identified the genes significantly upregulated in P organoids from multi-stroma co-cultures compared to PDAC organoids from multi-stroma co-cultures (hereon, ‘P organoid markers from multi-stroma co-cultures’), as well as the genes significantly upregulated in PDAC organoids from multi-stroma co-cultures compared to P organoids from multi-stroma co-cultures (hereon, ‘PDAC organoid markers from multi-stroma co-cultures’). For human comparisons, as done previously, we identified the respective orthologs of the above markers, which we defined as ‘P organoid orthologs from multi-stroma co-cultures’ and ‘PDAC organoid orthologs from multi-stroma co-cultures’. P organoid markers from multi-stroma co-cultures significantly overlapped with the murine pancreatitis Ductal 1 and Ductal 2 markers previously identified (*P <* 1.0e-135 and *P <* 1.3e-33, respectively; **Fig. 6E** and **4F**; **Supplementary Table 3**). On the other hand, PDAC organoid markers from multi-stroma co-cultures significantly overlapped with the murine PDAC malignant markers previously identified (*P <* 1.5e-116; **Fig. 6F** and **4I**; **Supplementary Table 3**). P and PDAC organoid orthologs from multi-stroma co-cultures also significantly overlapped with the previously identified human pancreatitis ductal and PDAC malignant orthologs, respectively (*P <* 2.5e-04 and *P <* 1.9e-40, respectively; **Extended Data Fig. 6P-Q** and **2M-N**; **Supplementary Table 3**).

Since both organoid co-cultures with PSCs and multi-stroma significantly captured gene expression profiles of *in vivo* pancreatitis and PDAC epithelial cells, we leveraged the transcriptomic datasets of mouse models and both co-culture models to pinpoint pancreatitis and PDAC epithelial cell markers shared across conditions. This cross-model analysis identified 177 pancreatitis ductal markers (e.g., *Sox9*, *Stat1*) and 254 PDAC malignant markers (e.g., *Muc5ac*) (**Supplementary Table 8**). Multiplex immunofluorescence (IF) analysis revealed a significantly higher abundance of MUC5AC^+^CK19^+^ epithelial cells in murine PDAC compared to pancreatitis tissues, normalized to CK19^+^ epithelial cells (**Fig. 6G-H**). In contrast, SOX9^+^CK19^+^ and STAT1^+^CK19^+^ epithelial cells were more abundant in pancreatitis tissues (**Fig. 6I-K**).

Altogether, our study shows how combining *in vitro* and *in vivo* models can help pinpoint epithelial cell markers that are differentially abundant in pancreatitis or PDAC tissues.

### Optimised multi-stromal organoid co-cultures better mimic fibroblast heterogeneity of pancreatitis and PDAC

After validating that P and PDAC organoid/multi-stroma co-cultures captured features of pancreatitis and PDAC epithelial cells *in vivo*, we set out to determine whether these co-cultures more closely mirror the transcriptome of fibroblasts *in vivo* compared to what we observed with PSC co-cultures. We leveraged the expression of *Gfp* and *TdTomato*, as well as fibroblast and mesothelial cell markers to identify PSCs, fibroblasts and mesothelial cells in the scRNA-seq dataset of P and PDAC organoid/multi-stroma co-cultures (**Fig. 6C** and **Extended Data Fig. 6M**). To evaluate differences in stromal composition of these multi-stroma co-cultures, we initially compared the transcriptome of multi-stroma co-cultured with PDAC organoids (hereon, ‘PDAC multi-stroma’) to multi-stroma co-cultured with P organoids (hereon, ‘P multi-stroma’). Notably, we found that myofibroblastic signatures (e.g., collagen formation, ECM organisation) were enriched in PDAC multi-stroma, which we had seen in murine and human PDAC CAFs *in vivo*, while it had not been observed in PDAC PSCs (**Fig. 7A**, **1E**, **2I** and **5D**; **Supplementary Table 7**).

**Figure 7.**
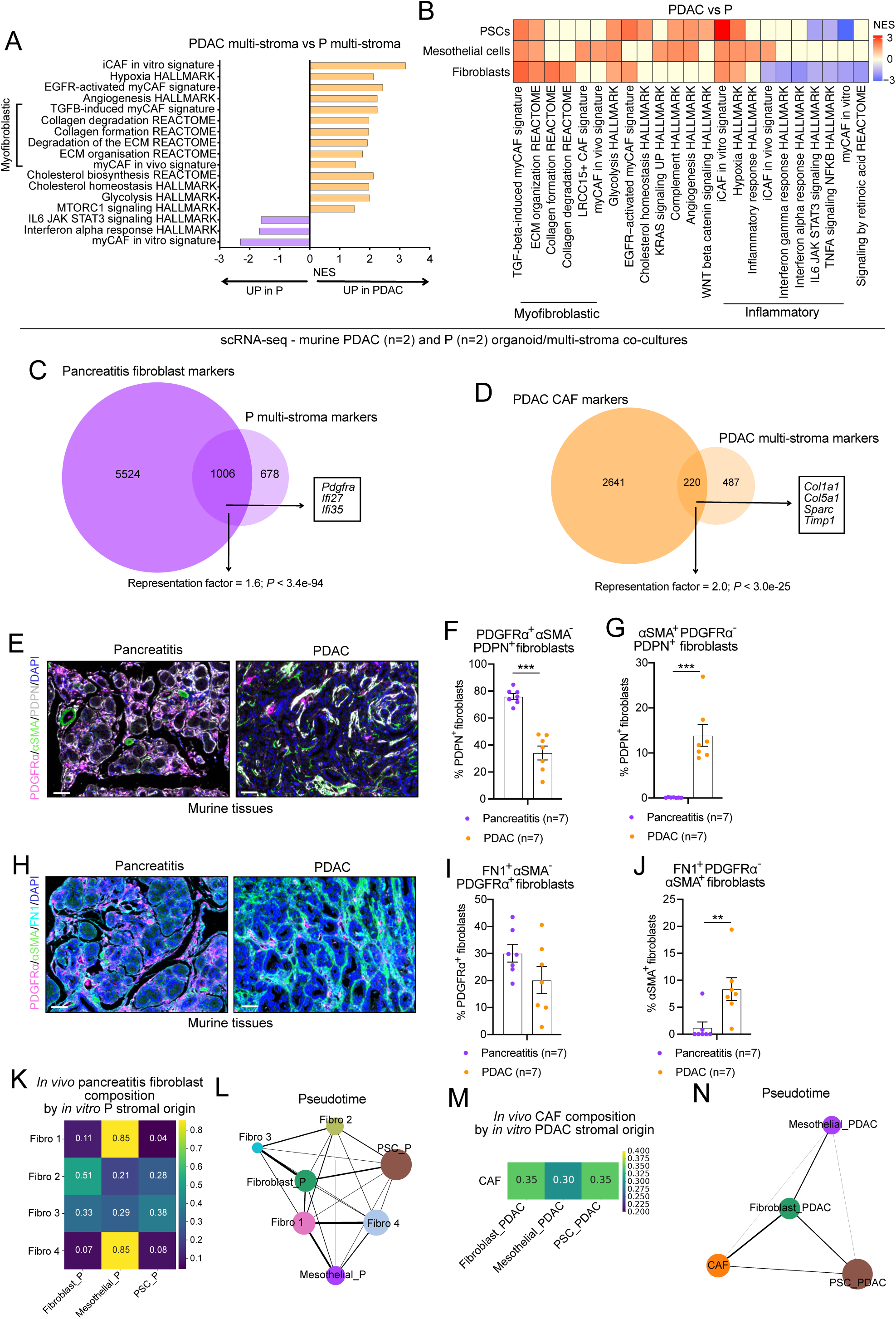
Optimised multi-stromal organoid co-cultures better mimic fibroblast heterogeneity of pancreatitis and PDAC. **(A)** Significantly upregulated and downregulated pathways identified by GSEA of murine multi-stroma (i.e. PSCs, fibroblasts and mesothelial cells) cultured with PDAC organoids compared to multi-stroma cultured with P organoids, as assessed by MAST from the scRNA-seq dataset of PDAC and P organoid/multi-stroma co-cultures. **(B)** Significantly upregulated and downregulated pathways identified by GSEA of PSCs, fibroblasts or mesothelial cells cultured with PDAC organoids compared to PSCs, fibroblasts or mesothelial cells cultured with P organoids, as assessed by MAST from the scRNA-seq dataset of PDAC and P organoid/multi-stroma co-cultures. **(C)** Venn diagrams of curated protein-coding genes significantly (*P* adj < 0.05) upregulated in multi-stroma cultured with P organoids compared to multi-stroma cultured with PDAC organoids (hereon, ‘P multi-stroma markers’), as assessed by scRNA-seq, and pancreatitis fibroblast markers. Significance of the overlap between datasets was defined by hypergeometric test (with denominator = 18,000). Selected genes common to both datasets are indicated. **(D)** Venn diagrams of significantly upregulated (*P* adj < 0.05) curated protein-coding DEGs in murine multi-stroma cultured with PDAC organoids compared to multi-stroma cultured with P organoids (hereon, ‘PDAC multi-stroma markers’), as assessed by scRNA-seq, and PDAC CAF markers. Significance of the overlap between datasets was defined by hypergeometric test (with denominator = 18,000). Selected genes common to both datasets are indicated. **(E)** Representative multiplex IF images of PDPN (white), αSMA (green), PDGFRα (purple) and DAPI (nuclear stain, blue) in murine pancreatitis and PDAC tissues. Scale bars = 50 μm. **(F-G)** Quantification of PDGFRα^+^αSMA^-^PDPN^+^DAPI^+^ **(F)** and αSMA^+^PDGFRα^-^PDPN^+^DAPI^+^ **(G)** CK19^-^ cells relative to PDPN^+^CK19^-^DAPI^+^ fibroblasts in murine pancreatitis and PDAC tissues, as assessed by multiplex IF analysis. Results show mean ± SEM. *** *P* < 0.001, Mann-Whitney test. **(H)** Representative multiplex IF images of FN1 (cyan), αSMA (green), PDGFRα (purple) and DAPI (nuclear stain, blue) in murine pancreatitis and PDAC tissues. Scale bars = 50 μm. **(I)** Quantification of FN1^+^αSMA^-^PDGFRα^+^DAPI^+^CK19^-^ cells relative to PDGFRα^+^CK19^-^DAPI^+^ fibroblasts in murine pancreatitis and PDAC tissues, as assessed by multiplex IF analysis. Results show mean ± SEM. No significant difference was found, as assessed by Mann-Whitney test. **(J)** Quantification of FN1^+^PDGFRα^-^αSMA^+^DAPI^+^CK19^-^cells relative to αSMA^+^CK19^-^DAPI^+^ fibroblasts in murine pancreatitis and PDAC tissues, as assessed by multiplex IF analysis. Results show mean ± SEM. ** *P* < 0.01, Mann-Whitney test. **(K)** Heatmap showing the fractional contribution of *in vitro* scRNA-seq PSCs, mesothelial cells and fibroblasts from P organoid/multi-stroma co-cultures (columns) to *in vivo* snRNA-seq pancreatitis fibroblast subtypes (rows), derived from averaged scANVI soft label probabilities. **(L)** PAGA illustrating the inferred pseudotemporal trajectory among *in vitro* P stromal cells and *in vivo* pancreatitis fibroblast subtypes. Nodes (i.e., circles) represent cell subsets, and their size is scaled to the number of cells per cluster. Edges (i.e., line) represent the strength of connectivity (i.e., thickness). **(M)** Heatmap showing the fractional contribution of *in vitro* scRNA-seq PSCs, mesothelial cells and fibroblasts from PDAC organoid/multi-stroma co-cultures (columns) to *in vivo* snRNA-seq PDAC CAFs (row), derived from averaged scANVI soft label probabilities. **(N)** PAGA illustrating the inferred pseudotemporal trajectory among *in vitro* PDAC stromal cells and *in vivo* PDAC CAFs. Nodes (i.e., circles) represent cell subsets, and their size is scaled to the number of cells per cluster. Edges (i.e., line) represent the strength of connectivity (i.e., thickness).

We then moved to understand whether pancreatitis and PDAC epithelial cells differentially reprogram stromal cells of distinct origins. Analysis of PSCs, fibroblasts and mesothelial cells revealed that PDAC organoids induced certain myofibroblastic (e.g., ECM organisation) and inflammatory (e.g., iCAF *in vitro*) signatures across all three stromal cell types (**Fig. 7B** and **Supplementary Table 7**). However, this analysis also highlighted differences in how PDAC and pancreatitis epithelial cells reprogrammed these distinct stromal cells. For example, fibroblasts and PSCs, but not mesothelial cells, downregulated additional inflammatory pathways (e.g., NF-κB and JAK/STAT signalling) when cultured with PDAC relative to P organoids (**Fig. 7B**). Furthermore, fibroblasts, but not PSCs or mesothelial cells, upregulated collagen-associated pathways and downregulated interferon responses when cultured with PDAC compared to P organoids (**Fig. 7B**). Of note, these fibroblast-specific changes were also observed when comparing PDAC CAFs to pancreatitis fibroblasts from murine tissues, suggesting that this cell type may be driving at least some of the gene signatures observed *in vivo* (**Fig. 2I**).

To further investigate whether P and PDAC organoid/multi-stroma co-cultures better capture gene markers of pancreatitis and PDAC fibroblasts in murine and human tissues, we leveraged a similar approach to our previous cross-model and cross-species comparisons. For murine comparisons, we identified the genes significantly upregulated in P multi-stroma compared to PDAC multi-stroma (hereon, ‘P multi-stroma markers’) and the genes significantly upregulated in PDAC multi-stroma compared to P multi-stroma (hereon, ‘PDAC multi-stroma markers’). We then overlapped these datasets with the previously identified pancreatitis fibroblast markers and PDAC CAF markers, respectively. This analysis revealed larger overlaps for both comparisons relative to what we observed between P and PDAC PSCs from co-cultures and the respective fibroblast populations from murine tissues. Specifically, 48% of P PSC markers (n=438/905; *P <* 1.3e-14; **Fig. 5F**) and 60% of P multi-stroma markers (n=1,006/1,684; *P <* 3.4e-94; **Fig. 7C** and **Supplementary Table 3**) were common to the pancreatitis fibroblast markers. Furthermore, while only 15% of PDAC PSC markers were common to PDAC CAF markers (n=174/1,155; *P* < 0.23; **Fig. 5G**), 31% of PDAC multi-stroma markers were common to PDAC CAF markers, a significant overlap (n=220/707; *P <* 3.0e-25; **Fig. 7D** and **Supplementary Table 3**). We then evaluated whether these organoid/multi-stroma co-cultures also better captured the gene profiles of human fibroblasts. To do this, we identified the orthologs of P multi-stroma markers (hereon, ‘P multi-stroma orthologs’) and PDAC multi-stroma markers (hereon, ‘PDAC multi-stroma orthologs’). The degree of overlap of P PSC orthologs (*P <* 1.2e-10) or P multi-stroma orthologs (*P <* 9.3e-11) with human pancreatitis fibroblast orthologs was similar (**Extended Data Fig. 7A and 5F**; **Supplementary Table 3**). However, contrary to PDAC PSC orthologs, PDAC multi-stroma orthologs significantly overlapped with human PDAC CAF orthologs (*P <* 5.9e-18; **Extended Data Fig. 7B and 5G**; **Supplementary Table 3**). Finally, we leveraged datasets from organoid/multi-stroma co-cultures and mouse models to identify fibroblast markers enriched in pancreatitis or PDAC for protein validation in tissues (e.g., *PDGFRA* and *ACTA2*, respectively; **Extended Data Fig. 7A-B**). In line with our immunohistochemical and transcriptomic analyses, multiplex IF revealed a significant increase in PDGFRα^+^αSMA^-^PDPN^+^ fibroblasts in pancreatitis and αSMA^+^PDGFRα^-^PDPN^+^ fibroblasts in PDAC, relative to PDPN^+^ fibroblasts (**Fig. 7E-G**). Furthermore, in line with *FN1* expression being higher in PDAC CAFs, while FN1^+^PDGFRα^+^αSMA^-^ fibroblasts showed no significant difference, FN1^+^αSMA^+^PDGFRα^-^ fibroblasts were significantly more abundant in PDAC compared to pancreatitis (**Fig. 7H-J** and **Extended Data Fig. 7B**). These findings indicate that organoid/multi-stroma co-cultures better reflect fibroblast heterogeneity observed *in vivo* than organoid/PSC co-cultures, and that combining different pancreatitis and PDAC models improves representation of disease complexity.

Better understanding of stroma evolution in disease progression may help identify new ways to prevent, diagnose and treat PDAC. Therefore, we leveraged P and PDAC organoid/multi-stroma scRNA-seq datasets to infer the cell(s) of origin of fibroblasts in snRNA-seq profiles of pancreatitis and PDAC murine tissues. To this end, we mapped the transcriptional profiles of *in vitro*–derived stromal populations (i.e., *PSCs*, *fibroblasts*, and *mesothelial cells*) to *in vivo* pancreatitis fibroblast or PDAC CAF references using a semi-supervised scANVI framework^26^. In pancreatitis, the *Wt1*^High^*Cd55*^High^ Fibro 1 and *Wt1*^High^*Dpp4*^High^ Fibro 4 subsets exhibited strong transcriptional overlap with *mesothelial cells* (85%), with only minor similarity to *fibroblasts* (7-11%) or *PSCs* (4-8%) (**Fig. 7K**). This is consistent with a mesothelial cell of origin for these two Fibro subsets. Probabilistic mapping also showed that the *Pdgfra*^High^*Pi16*^High^ Fibro 2 subset had the strongest overall similarity to *fibroblasts* (51%), while the *Fap*^High^ Fibro 3 subset had strong similarity to both *fibroblasts* (33%) and *PSCs* (38%; **Fig. 7K**). Pseudotemporal trajectory analysis further suggested a stronger connection between *fibroblasts* and the *Fap*^High^ Fibro 3 subset (**Fig. 7L**). In contrast, PDAC CAFs showed a more composite signature reflecting mixed stromal origins, and pseudotemporal trajectory analysis suggested a stronger connection between *fibroblasts* and PDAC CAFs (**Fig. 7M-N**).

In summary, P and PDAC organoid/multi-stroma co-culture models significantly capture *in vivo* gene expression patterns of pancreatitis and PDAC fibroblasts, and highlight the potential contribution of distinct stromal cell types to PDAC CAFs and pancreatitis fibroblasts from tissues.

## DISCUSSION

Our study revealed new aspects of fibroblast heterogeneity, as well as epithelial cell-fibroblast crosstalk, in pancreatitis and PDAC. First, we found that epithelial cell and fibroblast features are distinct between PDAC and pancreatitis, with myofibroblastic signatures being enriched in PDAC CAFs. Second, we found that multi-stromal organoid co-cultures better capture pancreatitis and PDAC fibroblast features compared to organoid co-cultures with only PSCs. Third, leveraging these models, we identified PDAC and pancreatitis epithelial cell-specific reprogramming of stromal cells of different origin, and inferred the *in vivo* cells of origin of PDAC CAFs and pancreatitis fibroblasts. Finally, we demonstrated that cross-species and cross-model analyses can help pinpoint markers upregulated in PDAC or pancreatitis.

Considering that chronic pancreatitis is a risk factor for PDAC, dissecting how fibroblasts and epithelial cells differently crosstalk in these diseases could guide the development of much needed preventative, diagnostic and therapeutic strategies for PDAC. Combining *in vitro* and *in vivo* models could reveal PDAC-specific features that may serve as early detection biomarkers of PDAC in chronic pancreatitis patients. They could also pinpoint candidate targets of fibroblast-epithelial cell crosstalk for the development of PDAC therapeutic strategies. Finally, they could help find targetable vulnerabilities of chronic pancreatitis, which remains incurable. Therapeutic interventions that are effective in impairing chronic pancreatitis could reduce the risk of developing PDAC in this patient group^42,43^. Although these long-term goals will require significant more work, the development of new models to investigate epithelial cell and fibroblast markers and crosstalk in pancreatitis and PDAC is a first step towards them. The organoid/multi-stroma co-culture models established in this study could also be leveraged in the future to shed light on pancreatitis and PDAC stromal cell-to-stromal cell crosstalk, which remains underexplored^3^.

While we leveraged caerulein-based models, as they are most frequently used to study pancreatitis in mice, future analyses of additional models, including GEMMs carrying mutations known to cause hereditary pancreatitis could complement our work^21,44–46^. Furthermore, in this study we focused on models treated with caerulein for six weeks as this prolonged treatment led to more fibrosis and ADM. However, shorter caerulein treatments for the generation of acute pancreatitis models have been shown to also alter the transcriptome of fibroblasts and epithelial cells^47^. Thus, it remains to be evaluated whether epithelial cell-fibroblast cross-talks are distinct in acute compared to prolonged pancreatitis. This may be relevant for the identification of phenotypic and functional changes specific to prolonged pancreatic inflammation, considering that only chronic, and not acute, pancreatitis increases the risk of developing PDAC in humans^48^. Finally, it remains to be determined how fibroblast-epithelial cell cross-talks evolve from inflammation to malignancy in the context of oncogenic drivers.

In PDAC, PDGFRα and αSMA are enriched in iCAFs and myCAFs, respectively^49^. Here, we find more PDGFRα^+^αSMA^-^ fibroblasts in pancreatitis and more αSMA^+^PDGFRα^-^ fibroblasts in PDAC. Consistent with this, pancreatitis fibroblasts show higher inflammatory signatures and fewer myofibroblastic features compared to PDAC CAFs. This observation suggests that myofibroblastic features are more specific to the malignant state. Further work will be required to determine whether targeting myCAF features in PDAC could be a therapeutic strategy. Indeed, while depletion of αSMA^+^ cells, which include myCAFs, or collagen I promoted PDAC progression^50,51^, selective targeting of myCAF subsets reduced PDAC metastases and therapy resistance^7,52^.

MyCAF and iCAF states that were initially identified in PDAC were later discovered also in other malignancies^53^. Thus, while our work focused on fibroblast heterogeneity in pancreatitis, it may help understand features of fibroblasts in other inflammatory states, and vice versa. Notably, fibroblast phenotypical and functional heterogeneity has been previously described in other inflammatory diseases^54–58^. For example, in rheumatoid arthritis, while *Fap* and *Pdpn* marked all fibroblasts, presence or absence of *Thy1* (encoding CD90) expression defined populations with distinct roles^54^. Our analyses suggest that *Thy1* and *Fap* are heterogeneously expressed in pancreatitis fibroblasts, while *Pdpn* expression is more uniform. Future genetic perturbation studies in mouse models may highlight shared phenotypical and functional features of fibroblast subsets across inflammatory conditions.

## MATERIALS AND METHODS

### Mouse models

Male and female C57BL/6J (strain number 632, Charles River, RRID:IMSR_JAX:000664) were purchased from the Charles River Laboratory (7-8-week-old at arrival). C57BL/6J-background KPC (*Kras*^LSL-G12D/+^; *Trp53*^R172H/+^; *Pdx1*-Cre) mice^22^ were bred in the Institute and palpated to follow tumour formation from two months of age. Presence of tumours was confirmed by ultrasound-based imaging. All animals were housed in accordance with the guidelines of the UK Home Office “Code of Practice for the Housing and Care of Experimental Animals”. They were kept behind strict barriered housing, which has maintained animals at a well-defined microbiological health status. This accommodation precludes access by wildlife, including rodent and insect vectors, and is free of infestation with ectoparasites. All animals were fed expanded rodent diet (Labdiet) and filtered water *ad libitum*. Environmental enrichment includes nesting material, structures for three-dimensional use of the cage and an area to retreat, and provision of chew blocks. All animal procedures and studies were reviewed by the Cancer Research UK Cambridge Institute (CRUK-CI) AWERB, approved by the Home Office and conducted under project license number PP4778090 in accordance with relevant institutional and national guidelines and regulations.

### Establishment of pancreatitis and acute pancreatitis mouse models

To model pancreatitis and acute pancreatitis, 8-9-week-old C57BL/6J mice were treated with the cholecystokinin analogue caerulein (C9026; Sigma-Aldrich) in endotoxin-free Dulbecco′s PBS (TMS-012-A, Sigma-Aldrich). Two caerulein-induced pancreatitis regimens were used in this paper. To model acute pancreatitis, mice were administered 100 μg/kg buprenorphine (57446/4003, Virbac) subcutaneously 1 hour before 6-8 hourly intraperitoneal injections of 75 μg/kg caerulein per day for two days (modified from Lugea et al.^59^). Forty-eight hours after the first caerulein dose of the second day of treatment, mice were sacrificed, and tissues harvested. To model pancreatitis, mice were treated with 250 μg/kg caerulein twice daily, as previously performed^43^, although we extended the treatment from two to six weeks based on our pilot data (not included) that showed longer treatment led to increased fibrosis. Gel cups were provided from week 4 to mitigate the body weight loss typically observed in pancreatitis mouse models. After six weeks of treatment, pancreatitis mice were sacrificed, and tissues harvested.

### Amylase and lipase activity plasma measurements

At experimental endpoints, mice were anesthetized with isoflurane, and blood was collected by cardiac puncture. Blood samples were collected in Multivette 600 K3E tubes containing EDTA (15.1671; Sarstedt) as an anticoagulant. Plasma was isolated from the blood by centrifugation at 10,000 rpm for 6 min at 4°C, and the plasma fraction was collected and stored at −80°C until analysis. Amylase and lipase activity levels in plasma were measured using the kits from Abcam (Ab102523 and Ab102524, respectively) following the manufacturer’s instructions.

### Establishment and culture of murine normal, pancreatitis and PDAC organoids

All PDAC organoids used in this study have undergone loss of heterozygosity (LOH) of the *Trp53* WT allele, apart from the KPC organoid line used in Extended Data Fig. 4 (see Oni and Biffi et al.^33^). Murine pancreatic intraepithelial neoplasia KC organoid line (C57BL/6J background) was previously described^33^. Murine PDAC KPC organoid lines (C57BL/6J background) T6-LOH and GB-T12-LOH were previously described^7,33^ apart from GB-T11-LOH and GB-T13-LOH. New murine normal pancreas (GB-N21-23, GB-N25 and GB-N27), P (GB-P1-5) and PDAC (GB-T11-LOH and GB-T13-LOH) organoids were established as previously described^7,29,33^. GB-T11-LOH and GB-T13-LOH were cultured with Nutlin-3a (SML0580; Sigma-Aldrich) from passage 0 as done previously for GB-T12-LOH organoids^7^. Briefly, to establish GB-T11-LOH and GB-T13-LOH organoids, C57BL/6J-background KPC tumours were digested with collagenase type XI (C9407, Sigma-Aldrich) and dispase (17105-041, Gibco). Bright field images of N, P and PDAC organoids were taken at 10X on day 1 post-isolation to assess and compare the morphological characteristics of different organoid lines. Organoids were propagated in complete media^29^. Organoid lines were typically cultured up to 30 passages, whenever possible, at 37°C with 5% CO2.

### Establishment and culture of murine PSCs, fibroblasts and mesothelial cells

Murine PSCs (SV40-immortalized, C57BL/6J background, PSC4 and PSC5) were previously described^4^. New murine pancreatic fibroblasts (GB-Fibro27 and GB-Fibro28) and pancreatic mesothelial cells (GB-PSC36-M-H11 and GB-PSC36-M-G1) were obtained while generating additional PSC lines from male and female C57BL/6J mice, respectively. Briefly, for each PSC line, two and a half pancreata from C57BL/6J mice were run on a density gradient centrifugation method with Histodenz (D2158; Sigma-Aldrich) and Gey’s Balanced Salt Solution (G9779; Sigma-Aldrich), as previously done^4^. While PSCs were established from the layer that forms in between the two interphases, fibroblasts were established from the pellets following centrifugation. Mesothelial cells were established by enriching for cobblestone-like cells found in fibroblast preparations by single-cell plating in 96-well plates. Both fibroblasts and mesothelial cells were immortalised using the pLVX-SV40 LT-IRES-tdTomato plasmid from Öhlund et al.^4^. Mouse PSCs, pancreatic fibroblasts and pancreatic mesothelial cells were cultured in DMEM (41966029; Gibco) containing 5% FBS. Cell lines were typically cultured up to 30 passages, whenever possible, at 37°C with 5% CO2. Cell line authentication was performed at the CRUK-CI. Mycoplasma testing for cell lines was performed prior to each freezing.

### Reverse transcription quantitative polymerase chain reaction analyses

RNA (1 μg) of PSCs, pancreatic fibroblasts, pancreatic mesothelial cells, as well as of KPC PDAC organoids and murine liver tissue (as controls), was reverse transcribed using TaqMan reverse transcription reagents (N808 0234; Applied Biosystems). qPCR was performed using gene-specific TaqMan probes (Thermo Fisher Scientific) and TaqMan master mix (4440040; Applied Biosystems). Gene expression was normalized to *Hprt*.

### Growth assays of organoids

For proliferation assays of N, P and PDAC organoids, 5,000 single cells were plated in 50 μL of 100% Matrigel on 24-well plates (Corning/Nunc) and cultured in 500 μL of complete media^29^. Organoid proliferation was followed for 120 hours with an Incucyte organoid module (Sartorius) with measurement of the organoid area per well every 3 hours (4 technical replicates per measurement). Data were normalised to the first measurement (i.e., at 0-hour post-plating on day 0).

### Western blot analyses

Organoids cultured in organoid complete media or reduced media were harvested in Cell Recovery Solution (354253; Corning) supplemented with protease inhibitors (11836170001; Roche) and phosphatase inhibitors (4906837001; Roche) and incubated for 30 min at 4°C. Cells were pelleted at 1500 *x g* for 5 min and lysed in 0.1% Triton X-100, 15 mmol/L NaCl, 0.5 mmol/L EDTA, 5 mmol/L Tris, pH 7.5, supplemented with protease inhibitors (11836170001; Roche) and phosphatase inhibitors (4906837001; Roche). Cells were incubated on ice for 30 min, briefly vortexed and pelleted at 13,200 rpm for 10 min at 4°C. Protein concentration of the collected supernatant was determined using the DC protein assay (5000113-5; Bio-Rad). Standard procedures were used for western blotting. Primary antibodies used were ACTIN (8456; Cell Signaling Technology; RRID:AB_10998774), p-STAT3 (9145; Cell Signaling Technology; RRID:AB_2491009), STAT3 (9139; Cell Signaling Technology; RRID:AB_331757), p16 (211542; Abcam; RRID:AB_2891084) and p19 (77184; Cell Signaling Technology), and SOX9 (AB5535; Merck; RRID: AB_2239761). Proteins were detected using appropriate HRP-conjugated secondary antibodies (Jackson ImmunoResearch Laboratories). All western blots are representative examples and have been repeated for at least two biological replicates.

### Bulk RNA-sequencing analyses of N, P and PDAC organoids in complete media

Bulk RNA-seq data of N, P and PDAC organoids cultured in complete media^29^ are available at the Gene Expression Omnibus (GEO) under the accession number GSE311286. Samples were collected in 1 mL of TRIzol Reagent (15596018; Invitrogen). RNA was extracted using the PureLink RNA mini kit (12183018A; Invitrogen). RNA concentration was measured using a Qubit and RNA quality was assessed on a TapeStation 4200 (Agilent) using the Agilent RNA ScreenTape kit. mRNA library preparations were performed using 55 μL of 10 ng/mL per sample (RNA integrity number, RIN ≥ 9). Illumina libraries were then sequenced on a NovaSeq6000. Differential gene expression and gene set enrichment analysis (GSEA) were performed as described previously^27^. GSEA significance was defined as NES >1.5 or <-1.5 and FDR < 0.25 as defined by Liberzon et al.^60^. Gene set variation analysis (GSVA) was performed using the GSVA R package^61^. Analyses are included in **Supplementary Table 4**.

#### Heatmap visualisation

Heatmaps were plotted using Morpheus (https://software.broadinstitute.org/morpheus).

### Immunohistochemical and histological analyses

Standard procedures were used for immunohistochemistry (IHC) analyses of tissues and P or PDAC organoid/PSC and multi-stroma co-cultures. Primary antibodies for IHC were αSMA (ab5694; Abcam; RRID:AB_2223021), PDPN (127403; BioLegend; RRID: AB_1134221), PDGFR-α (3174; Cell Signaling Technology; RRID: AB_2162345), CK19 (TROMA-III; DSHB; RRID: AB_2133570), Ly6G ( 127602; BioLegend; RRID: AB_1089180), and F4/80 (MCA497; Serotec; RRID: AB_323279). Hematoxylin (H-3404-100, Vector Lab) was used as nuclear counterstain. Masson’s trichrome and Hematoxylin & Eosin stains were performed according to standard protocols by the Histopathology core at the CRUK-CI. Briefly, following baking at 60°C for 1 hour, sections were dewaxed and rehydrated on Leica automated ST5020. The staining was performed on Leica’s automated Bond-III platform in conjunction with their BOND Polymer Refine Detection System (DS9800) and a modified version of their standard template. Antigen retrieval was performed at 100°C for 20 min in citrate or Tris-EDTA for F4/80 or PDGFRa, respectively. For aSMA, antigen retrieval was performed at 100°C for 10 min in Tris-EDTA. Antigen retrieval for CK-19 was performed by enzyme digestion (proteinase K) at 37°C using the Leica’s Bond enzyme concentrate (AR9551). Protein block (X090930-2; Dako) was applied for PDGFRa, CK-19, and F4/80. Primary antibodies used were listed above. The anti-rat secondary antibody (A110-322A; Bethyl Laboratories; RRID: AB_10681533) was applied for CK-19 and F4/80, and DAB Enhancer (AR9432; Leica) was used for all. Dehydration and clearing were performed on Leica automated ST5020 before sections were mounted on Leica CV5030 cover slipper. Stained sections were scanned with Aperio ScanScope CS and analysed using ImageScope software (RRID:SCR_014311) Positive Pixel Count algorithms or using HALO (RROD:SCR_018350), depending on the marker quantified. Images of tissue slides were obtained with an Axio Vert.A1 (ZEISS). The percentage of collagen area was determined by calculating the percentage of blue pixels relative to the entire stained area. To quantify αSMA stain, PDPN, PDGFR-α, CK19 and F4/80, the percentage of positive pixels was calculated relative to the entire stained section. To quantify Ly6G, the percentage of positive nuclei was calculated relative to the total number of nuclei. Stains and quantifications were performed blindly prior to plotting the data for visualization.

### Multiplex immunofluorescence analyses

Paraffin embedded tissue sections were dewaxed and rehydrated as per standard procedures then washed in deionized water. Tissue sections went through five rounds of antigen retrieval (AR), followed by blocking, incubation with primary antibody then secondary antibody and finally opal fluorophore. Briefly, AR was then performed by immersing the tissue slides in AR pH9 1X Tris-EDTA buffer (Abcam, ab93684) or AR pH6 1X AR6 buffer (Akoya Biosciences, AR600250ML) and boiling in a microwave for 20 min. After cooling, tissue slides were washed in deionized water by rocking twice for 2 min and then a further 2 min washing in PBS + 0.1% Tween20 (PBS-T). Tissue sections were then outlined using a hydrophobic barrier pen and blocking buffer (Akoya Biosciences, ARD1001EA) was incubated on the slides at room temperature (RT) for 15 min in a humidified chamber. Slides were incubated with primary antibody for 45 min at RT or 4°C overnight. Washing was performed by rocking the slides in PBS-T for 3 x 2 min. This was followed by incubation with secondary antibody (Akoya Biosciences, ARH1001EA) for 15 min at room temperature, washing, then incubation with opal fluorophore for 12 min. Slides were washed again then immersed in antigen retrieval buffer to being the cycle from microwaving to fluorophore incubation for all other primary antibodies in the panel. After the last round of staining incubation with opal fluorophore 780, DAPI was added to the tissues for 5 min. After washing, the slides were then mounted in ProLong Gold Antifade mountant (Thermofisher, P36930) and 1.5 mm coverslip (2980-245, Corning). The slides were then imaged using the Vectra Polaris. Tissue staining was analyzed using QuPath software. All sections were set to the same signal thresholds per channel, annotated by area to analyze and then cells detected using DAPI signal to identify each nuclei using the cell detection tool. Positivity for the markers of interest was defined by signal threshold in the object classifer tool and all parameters applied to each section remained the same. Primary antibodies used to identify these markers are as follows: αSMA (ab5694; Abcam; RRID:AB_2223021), PDPN (127403; BioLegend; RRID: AB_1134221), PDGFR-α (3174; Cell Signaling Technology; RRID: AB_2162345), FN1 (Ab2413; Abcam; RRID:AB_2262874), CK19 (TROMA-III; DSHB; RRID: AB_2133570), SOX9 (AB5535; Merck; RRID: AB_2239761), STAT1 (14994; Cell Signaling Technology; RRID:AB_2737027), MUC5AC (Ab3649, Abcam, RRID:AB_2146844). All buffers, anti-rabbit, anti-hamster and anti-mouse secondary antibodies, opal dyes and DAPI used were supplied in the Opal 6-Plex manual dection kit (Akoyabiosciences, NEL861001KT) unless otherwise detailed, additional goat-anti rat secondary (Vector, MP-7444-15) was used for antibodies raised in rat.

### Flow cytometry of murine PSCs, fibroblasts and mesothelial cells

Cells were cultured in 2D monolayer and then analysed by flow cytometry for CD200-APC (123809; Biolegend; RRID: AB_10900996) or treated with 100 ng/ml recombinant mouse interferon gamma (IFNγ, 315-05, PeproTech) for 48 hours and then analysed by flow cytometry for I-A/I-E-BV785 (MHCII, 107645; Biolegend; RRID: AB_2565977). IFNγ concentration was from Wijdeven et al.^62^.

### Flow cytometry and cell sorting of P and PDAC organoid/PSC co-cultures

PDAC or P-derived organoids were cultured alone or co-cultured with 35,000 of PCSs (per dome) per Matrigel done for 3.5 days in reduced media (i.e., 5% FBS DMEM)^4,7,27^ for flow cytometric and cell sorting analyses. Following single cell digestion of co-cultures, cells were blocked on ice with CD16/CD32 Pure 2.4G2 (553142, BD Bioscience; RRID: AB_394657) in PBS for 15 min.

For flow cytometric analysis of epithelial cells and subtypes of fibroblasts, cells were stained for 30 min on ice with anti-mouse CD326 (EpCAM)-AlexaFluor 488 (118210; BioLegend; RRID:AB_1134099), PDPN-APC/Cy7 (127418; BioLegend; RRID:AB_2629804), LIVE/DEAD Fixable Blue Dead Cell stain (L23105; Thermo Fisher Scientific). Then cells were washed in PBS twice, resuspend in PBS and analyzed on a CYTRK AURORA spectral flow cytometer. Flow analyses were performed blindly using FlowJo 10.8.2 (RRID:SCR_008520) prior to plotting the data for visualization.

Sorting of PDAC organoid/PSC co-cultures was performed following 3.5 days culture in reduced media (i.e. 5% FBS DMEM). Following single cell digestion of co-cultures, cells were stained for 30 min on ice with anti-mouse CD326 (EpCAM)-PE (118205; BioLegend; RRID:AB_1134176) and PDPN-AlexaFluor 488 (156208; BioLegend; RRID:AB_2814080).

Cells were resuspended in PBS with DAPI and sorted with a BD FACSMelody cell sorter.

### Bulk RNA-sequencing analyses of P organoids, PDAC organoids and PSCs flow-sorted from co-cultures and monocultures

Bulk RNA-seq data of P organoids, PDAC organoids and PSCs flow-sorted from monocultures and co-cultures are available at the GEO under the accession number GSE311286. Samples were collected in 1 mL of TRIzol Reagent (15596018; Invitrogen). RNA was extracted using the PureLink RNA mini kit (12183018A; Invitrogen). RNA concentration was measured using a Qubit and RNA quality was assessed on a TapeStation 4200 (Agilent) using the Agilent RNA ScreenTape kit. mRNA library preparations were performed using 55 μL of 10 ng/mL per sample (RIN ≥ 8.8). Illumina libraries were then sequenced on a NovaSeq6000. Differential gene expression and GSEA were performed as described previously^27^. GSEA significance was defined as NES >1.5 or <-1.5 and FDR < 0.25 as defined by Liberzon et al.^60^. GSVA was performed using the GSVA R package^61^. Analyses are included in **Supplementary Tables 5 and 6**.

#### NicheNet

NichenetR^63^ was applied on bulk RNA-seq data to infer the interaction between P or PDAC organoids and PSCs, following the approach described for PDAC organoid–PSC co-cultures in Lloyd et al.^27^.

#### Heatmap visualisation

Heatmaps were plotted using Morpheus (https://software.broadinstitute.org/morpheus).

### Single-cell RNA-sequencing of murine pancreatitis and PDAC organoid/multi-stroma co-cultures

PDAC or P-derived organoids were co-cultured with 30,000 of stromal cells (i.e., 10,000 each of PSCs, fibroblasts, and mesothelial cells) per Matrigel dome for 3.5 days in reduced media (i.e., 5% FBS DMEM)^4,7,27^. Green fluorescent protein (GFP)^+^ *Rosa26* KO PSCs, which carry the LRGN (LentisgRNA-EFS-GFP-neo) plasmid, were used for these experiments and were previously described^7^. Both fibroblasts and mesothelial cells are tdTomato^+^ as they were immortalised using the pLVX-SV40 LT-IRES-tdTomato plasmid from Öhlund et al.^4^. 10x Genomics 3’ V4 4-Plex (i.e., On-chip multiplexing, OCM) libraries were generated following the 3’ OCM protocol utilising standard and recommended reagents and conditions. Libraries were QC’ed via Qubit and TapeStation then pooled to 10 nM, loaded via qPCR targeting the P5/P7 adaptors and sequenced to a depth of ∼40,000 reads per cell on the NovaSeq X Plus device. Data is available at the GEO under the accession number GSE311279. Each OCM FASTQ file contains four samples. To demultiplex them into individual datasets, CellRanger “multi” function (10x Genomics) was used following the recommended workflow and a modified Mouse mm10 reference genome (2020-A (Jul 7, 2020)) that included reporter genes such as *Tdtomato* and *Gfp*. From the resulting output, the *raw_feature_bc_matrix* files were used as input for CellBender to remove ambient RNA, with an FDR of 0.01^64^. Each sample then underwent standard quality control using Scanpy^65^. Low-quality cells and outliers were excluded based on total counts (1st and 99th percentiles), gene counts (<100 genes), and mitochondrial gene content (>5%). Doublets were detected, and later removed, using the SOLO model implemented in scvi-tools^26,66^. Finally, all samples were concatenated into a single *Anndata* object and integrated using scvi-tools^26^. After integration, data was clustered using Leiden algorithm based on 2000 highly variable genes and top 30 PCs^67^. Then, cell clusters were annotated to their cell types based on known gene marker and reporter gene expression (*Gfp*: PSCs, *TdTomato*: fibroblasts and mesothelial cells). Analyses are included in **Supplementary Table 7**.

### Single-nuclei isolation and RNA-sequencing of murine pancreatitis and PDAC tissues

Single nuclei isolation and RNA-sequencing of pancreatitis and PDAC tissues was performed as previously described^7,25^, with some modifications. Specifically, for PDAC tissues, 35K nuclei were submitted, while for pancreatitis tissues, 70K were submitted. Data are available at the GEO under the accession number GSE311531. CellRanger count (10x Genomics) was used following the recommended workflow and Mouse mm10 reference genome (2020-A (Jul 7, 2020)). The same analysis steps as for the 10x OCM dataset were applied, from ambient RNA removal to clustering. Cell clusters were then annotated to cell types based on previously reported marker genes. To validate malignant cell identity, copy number variation (CNV) analysis was performed as done previously^27^ using a python implementation of inferCNV of the Trinity CTAT Project (https://github.com/broadinstitute/inferCNV) to estimate the copy number status in each cell type. Fibroblasts were used as reference key. Anndata (h5ad) file was then converted to Seurat object using sceasy R package (https://github.com/cellgeni/sceasy) to perform R based analysis. CellChat R package^68^ was used with the recommended setting to infer and visualize the differential cell-cell communication between pancreatitis and PDAC for common cell types. For differential expression analysis, Model-based Analysis of Single-cell Transcriptomics (MAST) method was used and sample id was included as a latent variable^69^. Analyses are included in **Supplementary Table 2**.

### Single-cell RNA-sequencing of human chronic pancreatitis and PDAC tissues

Raw FASTQ files of 4 human chronic pancreatitis (excluding the autoimmune sample) and 5 human PDAC (untreated) samples were obtained from Dimitrieva et al.^20^ (dbGaP Study Accession: phs003751.v1.p1). The sample metadata is in **Supplementary Table 1**. scRNA-seq analysis was performed same as done above but using reference genome GRCH38 (refdata-gex-GRCh38-2024-A).

### Mapping *in vitro* stromal populations to *in vivo* fibroblast and CAF subclusters

To map transcriptional correspondence between *in vitro* stromal cell populations and *in vivo* pancreatitis fibroblast or PDAC CAF states, we used a probabilistic framework based on scVI and scANVI, implemented via scArches^26,70^. A reference model was trained on *in vivo* fibroblast and CAF subclusters (Fibro 1–4, CAF) using 4,000 highly variable genes (latent dimension = 20) and extended to a scANVI classifier with subcluster labels and an “Unknown” placeholder to maintain label consistency. *In vitro* stromal cells (PSC_P/PDAC, Fibroblast_P/PDAC, Mesothelial cell_P/PDAC) were analysed separately for pancreatitis and PDAC conditions. Each query AnnData object was initialised as “Unknown” projected into the reference latent space via scArches, and fine-tuned for 20 epochs. For each *in vitro* cell, soft label probabilities indicated transcriptional similarity to *in vivo* fibroblast or CAF subtypes. Additionally, pancreatitis fibroblast subclusters were projected onto the CAF reference to assess potential state transitions. Cluster-level mean probabilities were summarised as normalised confusion-matrix heatmaps. All analyses were done on mac M1 processor and the accelerator was set to ‘mps’.

### Partition-based graph abstraction (PAGA)

Pseudotemporal trajectories of annotated cell types were inferred using partition-based graph abstraction (PAGA), implemented in Scanpy via the sc.tl.paga function^71^.

### Marker gene lists

Marker gene lists were identified for multiple cell clusters from the scRNA-seq and snRNA-seq datasets using the rank_genes_groups function in Scanpy and ‘wilcoxon’ as preferred method^65^. Genes with *p* < 0.05 and expression in at least 10% of cells within the target cluster were retained. Gene lists used are included in **Supplementary Table 2**. The resulting marker lists were subsequently used for GSEA on RNA-seq data. The AUCell method that is implemented in the decoupleR python library was used to enrich gene lists on snRNA-seq data^72,73^.

### Hypergeometric test and Jaccard index

Hypergeometric test, odd ratio and jaccard index were calculated for overlapping genes between each two gene lists using GeneOverlap R package and are listed in **Supplementary Table 3**. Additional hypergeometric test calculations with representation factor was performed with http://nemates.org/MA/progs/overlap_stats.html and the results are shown on the figures. The universe was set to 18,000 or 16,076 (for human orthologs) to represent the number of curated protein-coding genes and including *CDKN2A* and *CDKN2B* (https://bioconductor.org/packages/release/bioc/html/GeneOverlap.html).

### Statistical analysis

GraphPad Prism software (RRID:SCR_002798), customized R, python scripts and Jupyter notebooks were used for graphical representation of data. Statistical analysis was performed using non-parametric Mann-Whitney test or Kruskal-Wallis test, unpaired or paired Student’s *t* test, Jaccard index, and hypergeometric test (with a denominator = 18,000 for curated protein-coding genes for murine comparisons and 16,076 for non-redundant, mouse to human curated protein-coding orthologs for cross-species comparisons). All statistical details of experiments are specified in the figure legends and/or panel figures, including the number of technical and biological replicates, and how significance was defined.

## ACKNOWLEDGEMENTS

The authors would like to acknowledge the core facilities of the CRUK Cambridge Institute, and in particular Research Instrumentation and Cell Services, Biological Resource Unit, Histopathology, Genomics, Flow Cytometry, and IT and Scientific Computing. This work was largely supported by G.B. UK Research & Innovation Future Leaders Fellowship and G.B. Cancer Research UK Institutional core grant, with initial contributions from an Isaac Newton Trust/Wellcome ISSF/Uni of Cambridge Joint Grant. M.J., W.Lu. and S.H. were supported by a CRUK-NCI Cancer Grand Challenge grant (team CANCAN). The authors would also like to acknowledge Professor Howard Crawford (Henry Ford Health System) and Dr Tim Halim (Cancer Research UK Cambridge Institute) for advice on the caerulein treatment regimens.

## AUTHOR CONTRIBUTION

W.L. and M.J. conceptualised the work, performed the experiments and wrote the manuscript. G.M., M.Z., E.G.L., J.A.H., P.S.W.C., S.H., S.M., P.M.J., W.Lu., A.D., and R.B performed the experiments. P.K. co-supervised M.J with G.B. A.P. advised on sequencing analyses. M.V. supervised the mouse work. G.B. conceptualised and supervised the work, performed the experiments and wrote the manuscript.

## EXTENDED DATA FIGURES AND SUPPLEMENTARY TABLES

**Extended data Figure 1.**
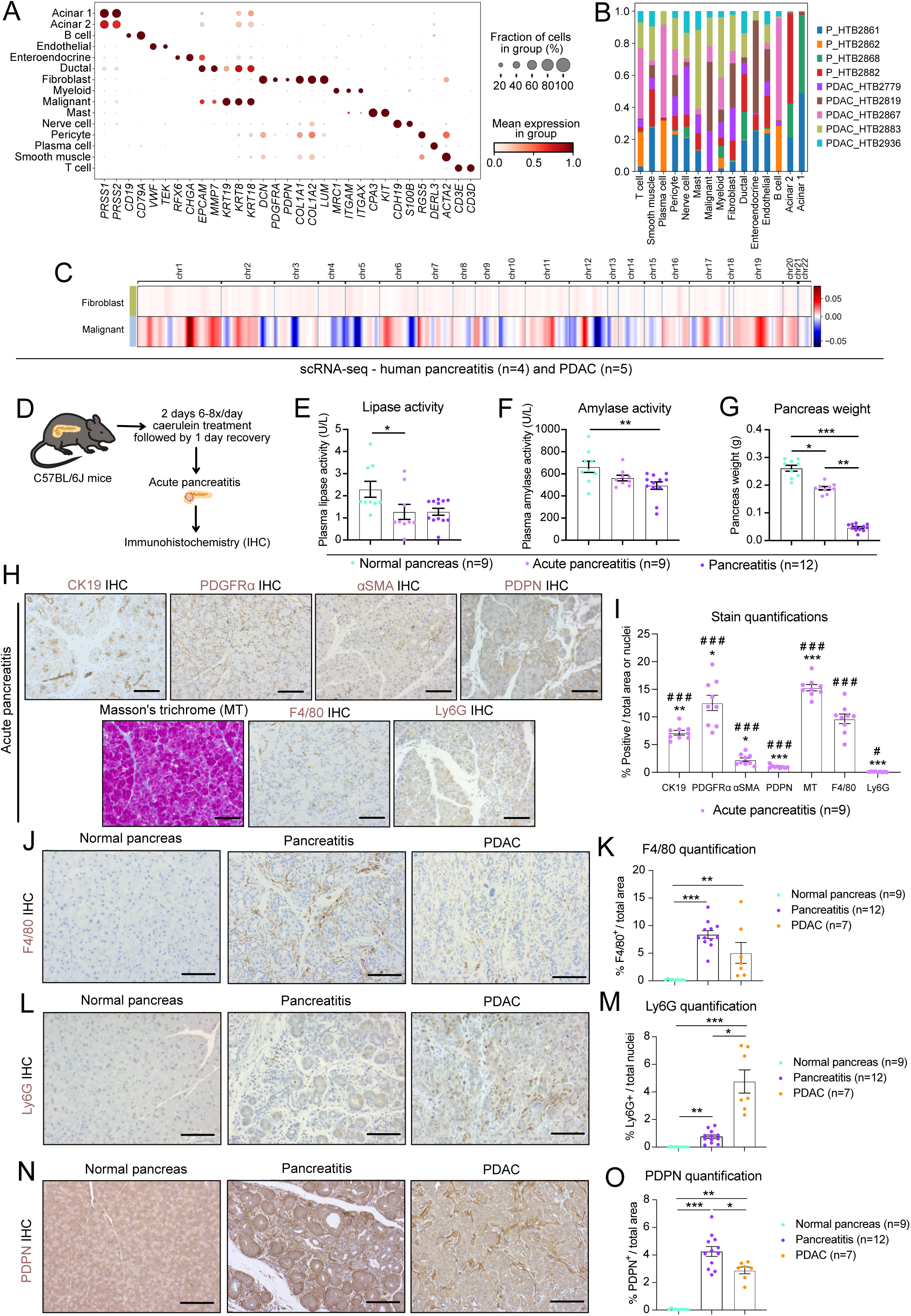
Myofibroblasts are more abundant in human and murine PDAC relative to pancreatitis. **(A)** Dot plot of scaled expression of cell type-specific markers in each cell cluster of human chronic pancreatitis (n=4) and pancreatic ductal adenocarcinoma (PDAC, n=5) tissues, as analysed by single-cell RNA-sequencing (scRNA-seq). The colour intensity represents the expression level, and the size of the dots represents the percentage of expressing cells. scRNA-seq analysis is from Dimitrieva et al. **(B)** Sample contribution to different cell types in human pancreatitis and PDAC analysed by scRNA-seq, represented as bar plot showing proportions of the different samples in each cell cluster. **(C)** Heatmap showing large-scale copy number variation (CNV) profile of the fibroblast and malignant cell clusters identified by scRNA-seq of human PDAC. The colour coding represents the CNV level based on a sliding window of 250 gene expression. Amplifications are shown in red, and deletions are shown in blue. The fibroblast cluster was used as reference cell cluster. **(D)** Schematic of models and techniques used for the analysis of murine acute pancreatitis tissues. **(E)** Lipase activity levels in plasma from normal pancreas, acute pancreatitis and pancreatitis mouse models. Results show mean ± SEM. * *P adj* < 0.05, Kruskal-Wallis test. **(F)** Amylase activity levels in plasma from normal pancreas, acute pancreatitis and pancreatitis mouse models. Results show mean ± SEM. ** *P adj* < 0.01, Kruskal-Wallis test. **(G)** Weights of pancreata from normal pancreas, acute pancreatitis and pancreatitis mouse models. Results show mean ± SEM. * *P adj* < 0.05; ** *P adj* < 0.01; *** *P adj* < 0.001, Kruskal-Wallis test. **(H-I)** Representative images and quantifications of the ductal marker keratin 19 (CK19), the fibroblast marker platelet-derived growth factor receptor alpha (PDGFRα), the myofibroblast marker alpha smooth muscle actin (αSMA), the fibroblast marker podoplanin (PDPN), the macrophage marker F4/80, the neutrophil marker Ly6G and Masson’s trichrome (for collagen deposition) stains of acute pancreatitis murine tissues. Results show mean ± SEM. * *P* < 0.05; ** *P* < 0.01; *** *P* < 0.001, Mann-Whitney test, indicate significant downregulation compared to pancreatitis tissues from Figures 1 and S1. ^#^ *P* < 0.05; ^###^ *P* < 0.001, Mann-Whitney test, indicate significant upregulation compared to normal pancreas tissues from Figures 1 and S1. Scale bars, 50 μm. **(J-M)** Representative images and quantifications of F4/80 **(J-K)**, Ly6G **(L-M)** and PDPN **(N-O)** stains of normal pancreas, pancreatitis and PDAC murine tissues. Results show mean ± SEM. * *P adj* < 0.05; ** *P adj* < 0.01; *** *P adj* < 0.001, Kruskal-Wallis test. Scale bars, 50 μm.

**Extended data Figure 2.**
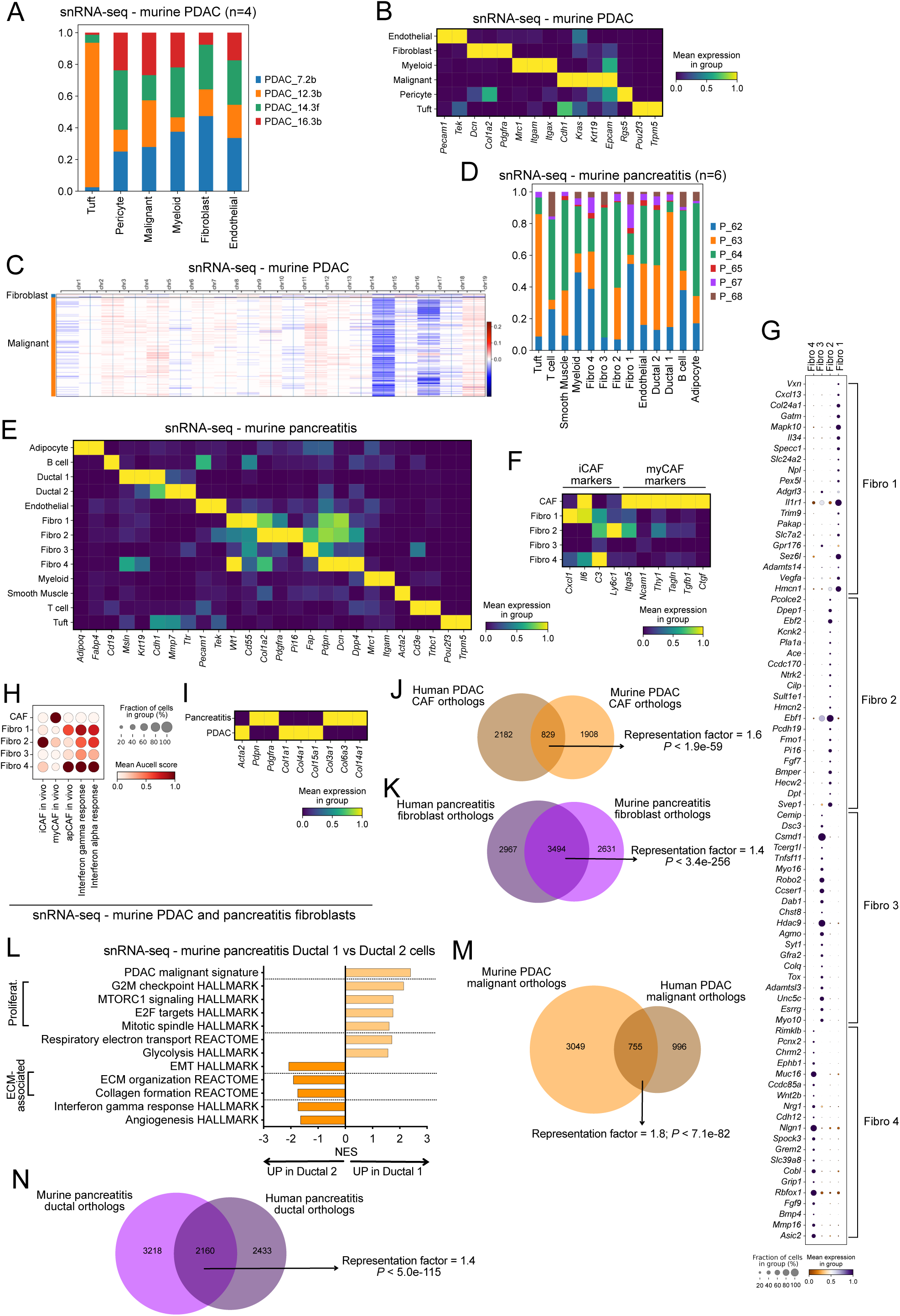
Fibroblast and epithelial cell populations are more heterogeneous in pancreatitis than PDAC mouse models. **(A)** Sample contribution to different cell types in murine PDAC (n=4), represented as bar plot showing proportions of the different tumour samples in each cell cluster, as analysed by single-nuclei RNA-sequencing (snRNA-seq). **(B)** Heatmap of scaled expression of cell type-specific markers in each cell cluster of murine PDAC. Data are scaled such that the cluster with the lowest average expression = 0 and the highest = 1 for each gene. **(C)** Heatmap showing large-scale CNV profile of the fibroblast and epithelial malignant cell clusters identified by snRNA-seq of murine PDAC. The colour coding represents the CNV level based on a sliding window of 250 gene expression. Amplifications are shown in red, and deletions are shown in blue. The fibroblast cluster was used as reference cell cluster. **(D)** Sample contribution to different cell types in murine pancreatitis tissues (n=6), represented as bar plot showing proportions of the different samples in each cell cluster, as analysed by snRNA-seq. **(E)** Heatmap of scaled expression of cell type-specific markers in each cell cluster of murine pancreatitis. Data are scaled such that the cluster with the lowest average expression = 0 and the highest = 1 for each gene. **(F)** Heatmap of scaled expression of iCAF (*Cxcl1*, *Il6*, *C3*, *Ly6c1*) and myCAF (*Itga5*, *Ncam1*, *Thy1*, *Tagln*, *Tgfb1*, *Ctgf*) markers in each pancreatitis and PDAC fibroblast cluster. Data are scaled such that the cluster with the lowest average expression = 0 and the highest = 1 for each gene. **(G)** Dot plot of scaled expression of the top 20 markers for each pancreatitis fibroblast cluster compared to the other pancreatitis fibroblast clusters (expression percentage cut-off 10%, *P adj* < 0.05, Wilcoxon test). The colour intensity represents the expression level, and the size of the dots represents the percentage of expressing cells. **(H)** Dot plot of scaled Aucell score enrichment of selected signatures in each fibroblast cluster from murine pancreatitis and PDAC snRNA-seq dataset. *In vivo* CAF signatures are from Elyada et al. Interferon signatures are from HALLMARK. **(I)** Heatmap of scaled expression of fibroblast markers (*Acta2*, *Pdpn*, *Pdgfra*) and collagens (*Col1a1*, *Col4a1*, *Col15a1*, *Col3a1*, *Col6a3*, *Col14a1*) in fibroblasts from murine pancreatitis and PDAC from the snRNA-seq dataset. Data are scaled such that the cluster with the lowest average expression = 0 and the highest = 1 for each gene. **(J)** Venn diagrams of significantly upregulated (*P* adj < 0.05) human ortholog curated protein-coding differentially expressed genes (DEGs) in murine PDAC CAFs compared to murine pancreatitis fibroblasts *in vivo* (hereon, ‘murine PDAC CAF orthologs’) and human PDAC CAFs compared to human pancreatitis fibroblasts *in vivo* (hereon, ‘human PDAC CAF orthologs’), as assessed by snRNA-seq and scRNA-seq, respectively. Significance of the overlap between datasets was defined by hypergeometric test (with denominator = 16,076 orthologs). **(K)** Venn diagrams of ortholog curated protein-coding DEGs significantly upregulated (*P* adj < 0.05) in murine pancreatitis fibroblasts compared to murine PDAC CAFs (hereon, ‘murine pancreatitis fibroblast orthologs’) and human pancreatitis fibroblasts compared to human PDAC CAFs (hereon, ‘human pancreatitis fibroblast orthologs’), as assessed by snRNA-seq and scRNA-seq, respectively. Significance of the overlap between datasets was defined by hypergeometric test (with denominator = 16,076 orthologs). **(L)** Significantly upregulated and downregulated pathways identified by Gene Set Enrichment Analysis (GSEA) of pancreatitis Ductal 1 compared to Ductal 2 cells, as assessed by Model-based Analysis of Single-cell Transcriptomics (MAST) from the snRNA-seq dataset, and genes were ranked based on log2 fold change. The PDAC malignant cell signature was obtained by comparing murine PDAC malignant cells to pancreatitis ductal cells from the snRNA-seq dataset. **(M)** Venn diagrams of differentially expressed curated protein-coding ortholog genes significantly (*P* adj < 0.05) upregulated in murine PDAC malignant cells compared to pancreatitis ductal cells (hereon, ‘murine PDAC malignant orthologs’) and human PDAC malignant cells compared to pancreatitis ductal cells (hereon, ‘human PDAC malignant orthologs’), as assessed by snRNA-seq and scRNA-seq, respectively. Significance of the overlap between datasets was defined by hypergeometric test (with denominator = 16,076 orthologs). **(N)** Venn diagrams of differentially expressed curated protein-coding ortholog genes significantly (*P* adj < 0.05) upregulated in murine pancreatitis ductal cells compared to PDAC malignant cells (hereon, ‘murine pancreatitis ductal orthologs’) and human pancreatitis ductal cells compared to PDAC malignant cells (hereon, ‘human pancreatitis ductal orthologs’), as assessed by snRNA-seq and scRNA-seq, respectively. Significance of the overlap between datasets was defined by hypergeometric test (with denominator = 16,076 orthologs).

**Extended data Figure 3.**
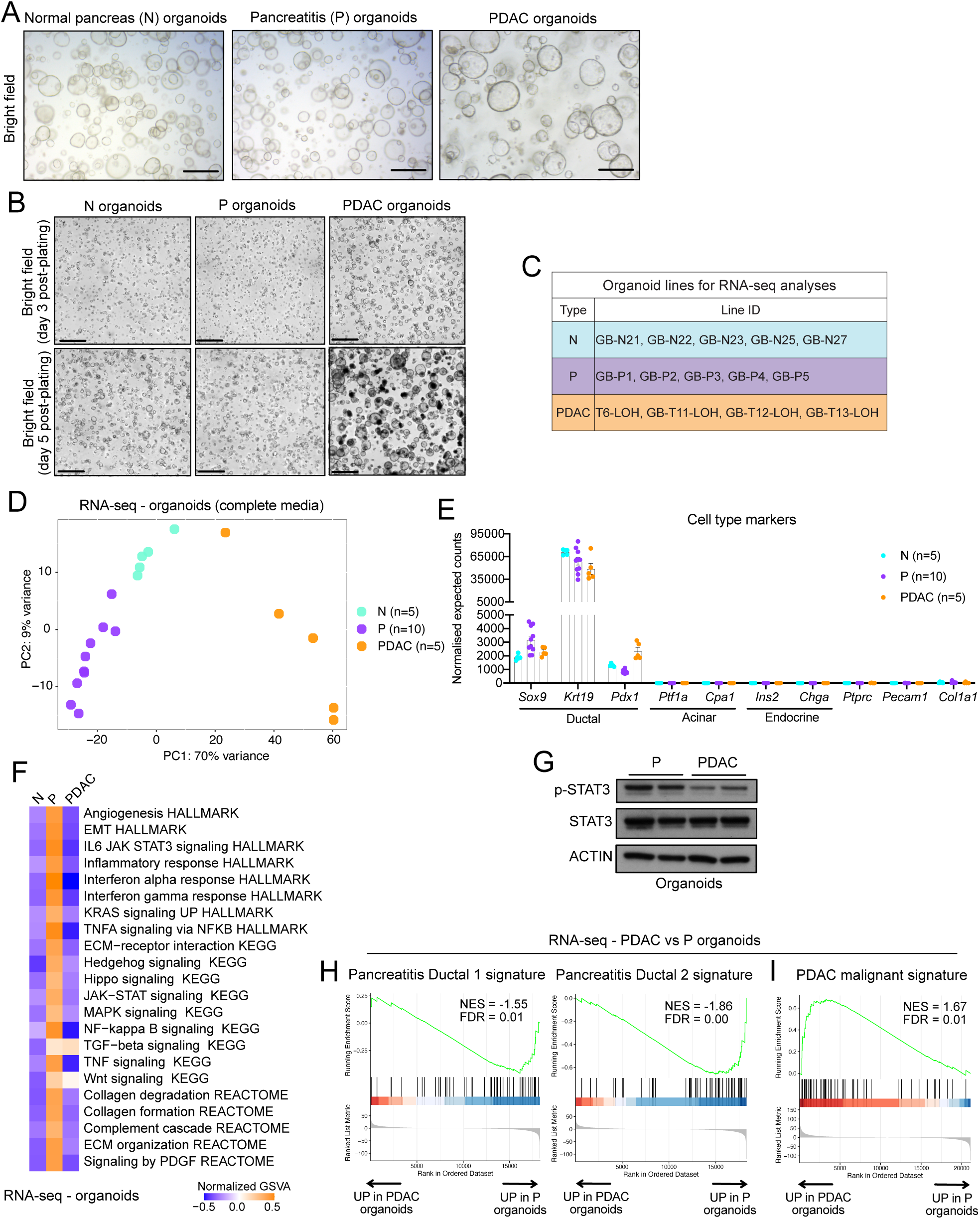
Pancreatitis-derived epithelial organoids capture *in vivo* gene expression patterns of pancreatitis ductal cells. **(A)** Bright field images of organoids generated from normal pancreas (hereon, termed N), pancreatitis (hereon, termed P) and PDAC tissues after > 5 passages; scale bars = 50 μm. **(B)** Bright field images (taken with an Incucyte organoid module) of N, P and PDAC organoids at day 3 and day 5 post-plating, as part of a growth assay; scale bars = 800 μm. **(C)** Summary table of the N, P and PDAC organoid lines analysed by RNA-sequencing (RNA-seq) after 3 days of culture in complete media. **(D)** Principal component analysis (PCA) of N (n=5 biological replicates), P (n=5 biological replicates, each with 2 technical replicates at different passages) and PDAC (n=4 biological replicates, of which one with 2 technical replicates at different passages) organoids, as assessed by RNA-seq. **(E)** RNA-seq expression of ductal (*Sox9*, *Krt19*, *Pdx1*), acinar (*Ptf1a*, *Cpa1*), endocrine (*Ins2*, *Chga*), immune (*Ptprc*), endothelial (*Pecam1*) and fibroblast (*Col1a1*) markers in N, P and PDAC organoids. Results show mean ± SEM. **(F)** Heatmap showing Gene Set Variation Analysis enrichment (GSVA) scores (averaged per group) of selected pathways in N, P and PDAC organoids, as assessed by RNA-seq. Pathways displayed correspond to pathways significantly upregulated in GSEA in P organoids (Figures 3E and/or 3F). **(G)** Western blot of phospho-STAT3 (p-STAT3) and STAT3 in P and PDAC organoids cultured in complete media for 3 days. ACTIN, loading control. **(H)** GSEA of the murine pancreatitis Ductal 1 (left) and Ductal 2 (right) signatures in PDAC organoids compared to P organoids. The signatures were significantly downregulated in PDAC organoids (i.e. significantly enriched in P organoids). The signatures were obtained by comparing pancreatitis Ductal 1 and Ductal 2 cells, respectively, to PDAC malignant cells from the murine snRNA-seq dataset. NES, normalized enrichment score. FDR, false discovery rate. **(I)** GSEA of the PDAC malignant signature in PDAC organoids compared to P organoids. The signature was obtained by comparing PDAC malignant cells to pancreatitis ductal cells from the murine snRNA-seq dataset. The signature was significantly enriched in PDAC organoids.

**Extended data Figure 4.**
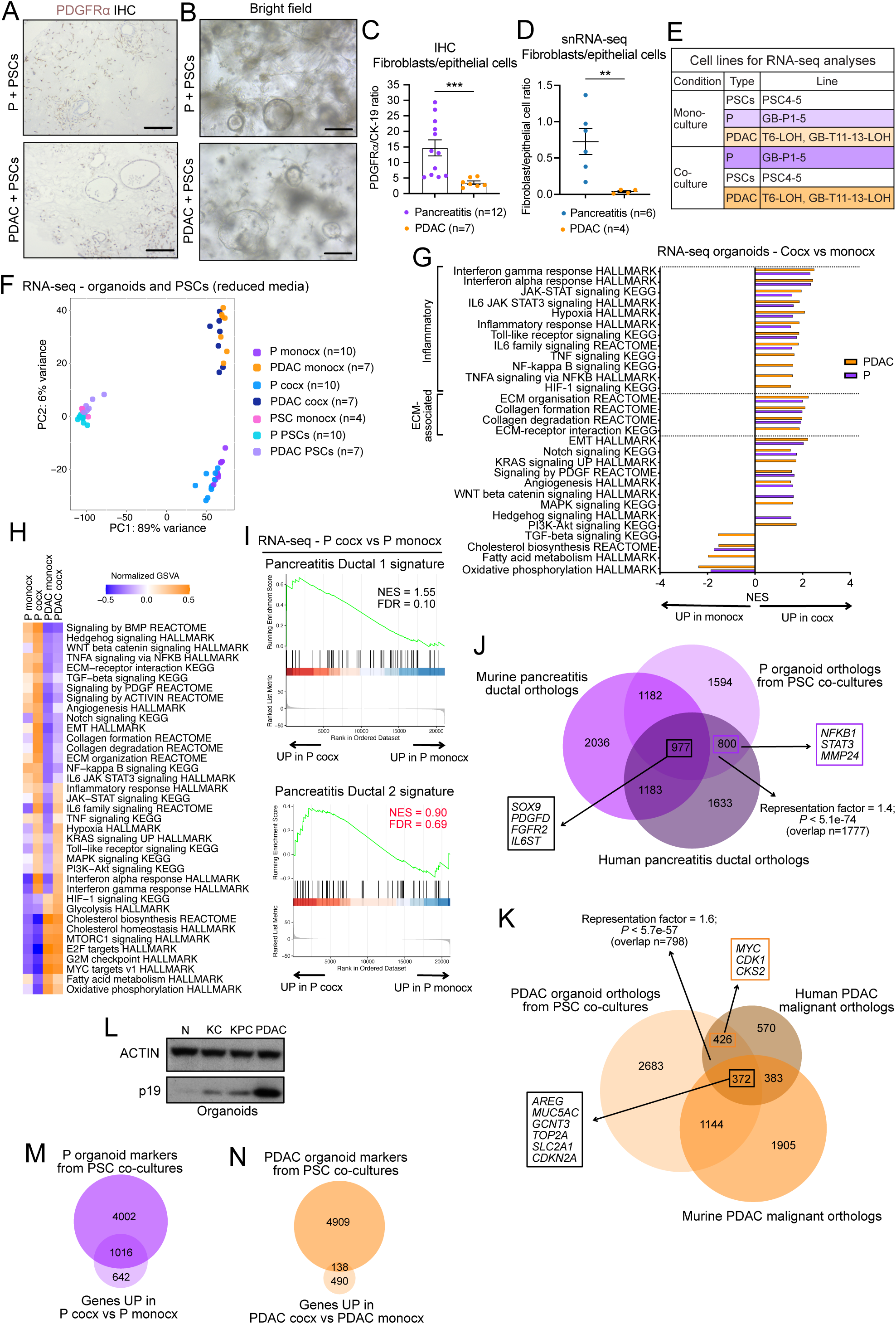
The transcriptome of pancreatitis ductal cells is more influenced by fibroblast signalling compared to PDAC malignant cells. **(A)** Representative images of PDGFRα stains of P and PDAC organoid/pancreatic stellate cell (PSC) co-cultures after 7 days in culture. Scale bars = 100 μm. **(B)** Bright field images of P and PDAC organoid/PSC co-cultures after 7 days in culture. Scale bars = 100 μm. **(C)** Fibroblast (PDGFRα^+^)/epithelial cell (CK19^+^) ratio as assessed by immunohistochemical analysis of pancreatitis or PDAC murine tissues (from Figure 1). Results show mean ± SEM. *** *P* < 0.001, Mann-Whitney test. **(D)** Ratio of fibroblast to epithelial cell numbers per sample as determined by snRNA-seq analysis of pancreatitis and PDAC murine tissues (from Figure 2). Results show mean ± SEM. ** *P* < 0.01, Mann-Whitney test. **(E)** Summary table of the conditions analysed by RNA-seq in monoculture (i.e. culture of a single cell type) or co-culture (i.e. culture of two or more cell types). **(F)** PCA of P organoids in monocultures (i.e., P monocx, n=5 biological replicates, each with 2 replicates of different passages) or in co-culture with PSCs (i.e., P cocx, n=10 biological replicates), PDAC organoids in monocultures (i.e., PDAC monocx, n=4 biological replicates, of which three samples with 2 replicates of different passages) or in co-culture with PSCs (i.e., PDAC cocx, n=7 biological replicates), and PSC monocultures (i.e., PSC monocx, n=2 biological replicates, each with 2 replicates of different passages) and in co-culture with PDAC organoids (i.e., PDAC PSCs, n=7 biological replicates) or P organoids (i.e., P PSCs, n=10 biological replicates), as assessed by RNA-seq. **(G)** Significantly upregulated and downregulated pathways identified by GSEA of P or PDAC organoids in co-culture compared to their respective monocultures, as assessed by RNA-seq. **(H)** Heatmap showing GSVA enrichment scores (averaged per group) of selected pathways in P and PDAC organoid monocultures and co-cultures, as assessed by RNA-seq. Pathways displayed correspond to pathways significantly altered in GSEA (Figures 4E and/or S4G). **(I)** GSEA of the murine pancreatitis Ductal 1 (top) and Ductal 2 (bottom) signatures in P organoids in co-culture with PSCs compared to P organoids in monoculture. The pancreatitis Ductal 1 signature was significantly enriched in P organoids in co-culture. The pancreatitis Ductal 2 signature was not significantly different between conditions. **(J)** Venn diagrams of differentially expressed curated protein-coding ortholog genes significantly (*P* adj < 0.05) upregulated in P organoids in PSC co-cultures compared to PDAC organoids in PSC co-cultures (hereon, ‘P organoid orthologs from PSC co-cultures’), as assessed by RNA-seq, murine pancreatitis ductal orthologs, and human pancreatitis ductal orthologs. Selected genes common to all or two datasets are indicated. Significance of the overlap between P organoid orthologs from PSC co-cultures and human pancreatitis ductal orthologs was defined by hypergeometric test (with denominator = 16,076 orthologs). **(K)** Venn diagrams of differentially expressed curated protein-coding ortholog genes significantly (*P* adj < 0.05) upregulated PDAC organoids in PSC co-cultures compared to P organoids in PSC co-cultures (hereon, ‘PDAC organoid orthologs from PSC co-cultures’), as assessed by RNA-seq, murine PDAC malignant orthologs, and human PDAC malignant orthologs. Selected genes common to all or two datasets are indicated. Significance of the overlap between PDAC organoid orthologs from PSC co-cultures and human PDAC malignant orthologs was defined by hypergeometric test (with denominator = 16,076 orthologs). **(L)** Western blot of p19 in N and PDAC organoids, as well as PDAC organoids that have not undergone loss of heterozygosity of the wild-type *Trp53* allele (KPC organoids) and pre-cancerous pancreatic intraepithelial neoplasia organoids from KC mice (KC organoids) cultured in complete media for 3 days. ACTIN, loading control. **(M)** Venn diagrams of curated protein-coding significantly (*P* adj < 0.05) upregulated genes in P organoids in co-culture with PSCs compared to PDAC organoids in co-culture with PSCs (hereon, ‘P organoid markers from PSC co-cultures’) and curated protein-coding significantly upregulated genes in P organoids in co-culture with PSCs compared to P organoids in monoculture. **(N)** Venn diagrams of curated protein-coding significantly (*P* adj < 0.05) upregulated genes in PDAC organoids in co-culture with PSCs compared to P organoids in co-culture with PSCs (hereon, ‘PDAC organoid markers from PSC co-cultures’) and curated protein-coding significantly upregulated genes in PDAC organoids in co-culture with PSCs compared to PDAC organoids in monoculture.

**Extended data Figure 5.**
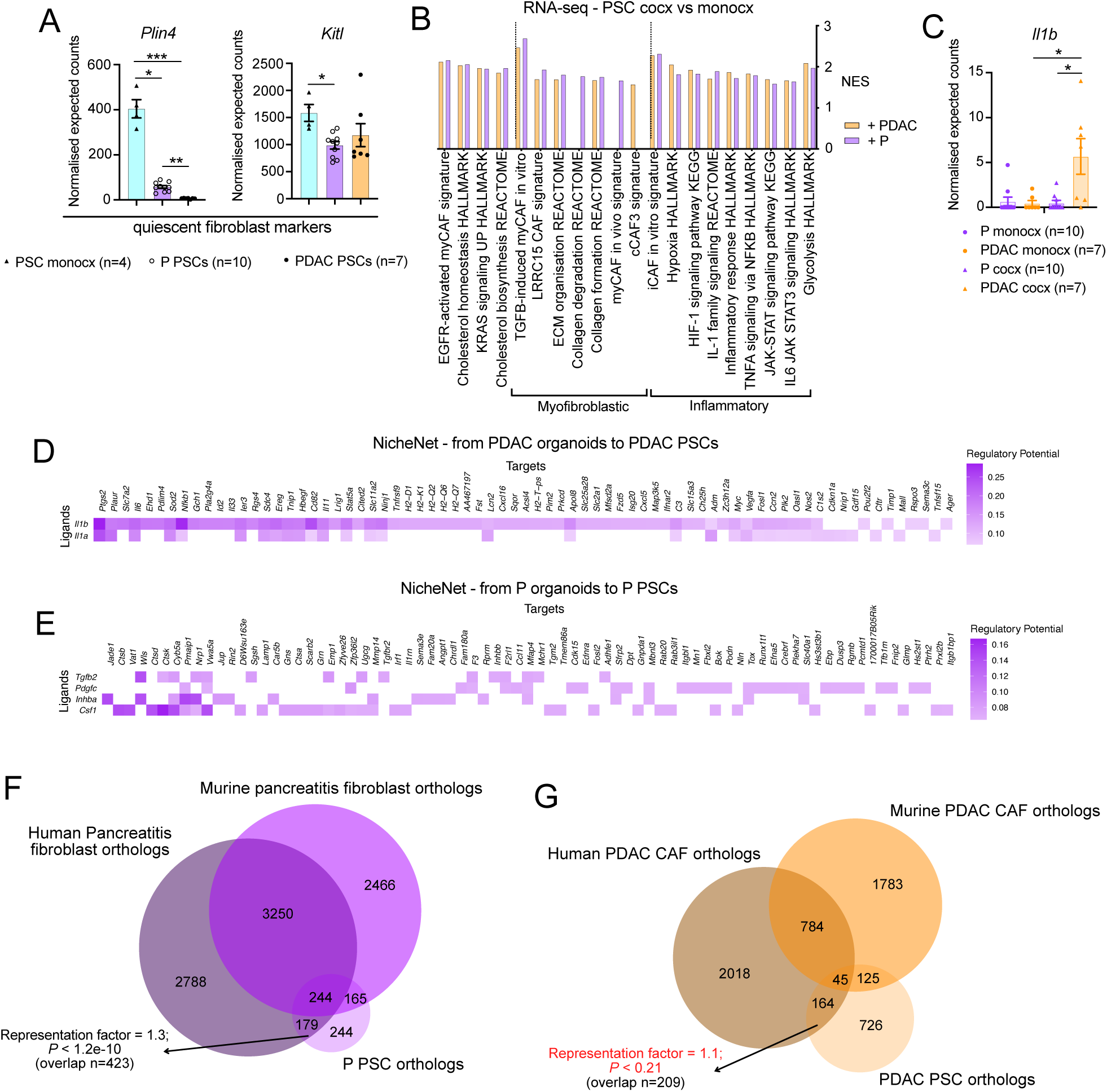
Organoid/PSC co-cultures only partially recapitulate the *in vivo* transcriptomic differences of pancreatitis and PDAC fibroblasts. **(A)** RNA-seq expression of quiescent fibroblast markers (*Plin4*, *Kitl*) in PSC monocx (n=2, 2 different passages) and co-cultures with P (n=10) or PDAC (n=7) organoids. Results show mean ± SEM. * *P adj* < 0.05; ** *P adj* < 0.01; *** *P adj* < 0.001, Kruskal-Wallis test. **(B)** Significantly upregulated pathways identified by GSEA of PSCs cultured with PDAC or P organoids compared to PSC monocx, as assessed by RNA-seq. **(C)** RNA-seq expression of *Il1b* in P or PDAC monocx and P or PDAC cocx with PSCs. Results show mean ± SEM. * *P adj* < 0.05, Kruskal-Wallis test. **(D)** Ligand–target heatmap shows top selected ligands of PDAC organoids inferred to regulate target genes in co-cultured PSCs, as assessed by NicheNet analysis. **(E)** Ligand–target heatmap shows top selected ligands of P organoids inferred to regulate target genes in co-cultured PSCs, as assessed by NicheNet analysis. **(F)** Venn diagrams of differentially expressed curated protein-coding ortholog genes significantly (*P* adj < 0.05) upregulated in PSCs cultured with P organoids compared to PSCs cultured with PDAC organoids (hereon, ‘P PSC orthologs’), as assessed by RNA-seq, murine pancreatitis fibroblast orthologs, and human pancreatitis fibroblast orthologs. Significance of the overlap between datasets was defined by hypergeometric test (with denominator = 16,076 orthologs). **(G)** Venn diagrams of differentially expressed curated protein-coding ortholog genes significantly (*P* adj < 0.05) upregulated in murine PSCs cultured with PDAC organoids compared to murine PSCs cultured with P organoids (hereon, ‘PDAC PSC orthologs’), as assessed by RNA-seq, murine PDAC CAF orthologs, and human PDAC CAF orthologs. Significance of the overlap between datasets was defined by hypergeometric test (with denominator = 16,076 orthologs).

**Extended data Figure 6.**
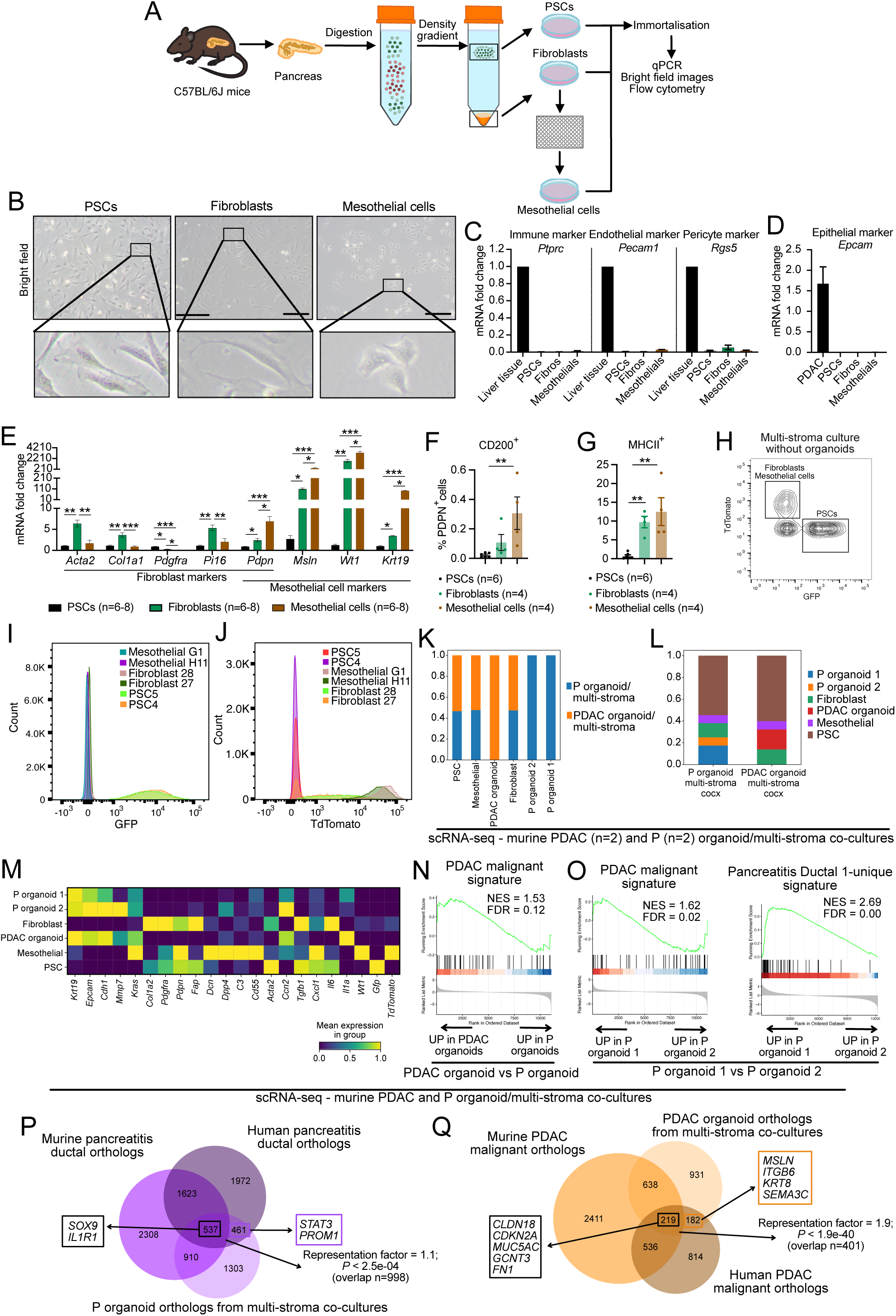
Cross-model analysis pinpoints epithelial cell markers of pancreatitis and PDAC. **(A)** Schematic illustrating how pancreatic fibroblast and mesothelial cell lines were generated and validated. **(B)** Bright field images of murine PSCs (left), fibroblasts (middle) and mesothelial cells (right) cultured in 2D. Scale bars = 100 μm. Bottom: Inserts show magnifications. **(C)** RNA expression levels of immune cell (*Ptprc*), endothelial cell (*Pecam1*) and pericyte (*Rgs5*) markers in PSCs (n=2, 2-3 technical replicates), fibroblasts (i.e., Fibros, n=2, 2-3 technical replicates) and mesothelial cells (i.e., mesothelials, n=2, 2-3 technical replicates) cultured in 2D, and in liver tissue (n=2 technical replicates). **(D)** RNA expression levels of epithelial cell marker (*Epcam*) in PSCs (n=2, 2-3 technical replicates), fibroblasts (i.e., Fibros, n=2, 2-3 technical replicates) and mesothelial cells (i.e., mesothelials, n=2, 2-3 technical replicates) cultured in 2D, and in PDAC organoids (i.e., PDAC, n=3). **(E)** RNA expression levels of fibroblast (*Acta2*, *Col1a1*, *Pdgfra*, *Pi16, Pdpn*) and mesothelial cell (*Pdpn*, *Msln*, *Wt1*, *Krt19*) markers in PSCs, fibroblasts and mesothelial cells cultured in 2D. Results show mean ± SEM of n=2 biological replicates with n=3-4 technical replicates per stromal cell line. * *P adj* < 0.05; ** *P adj* < 0.01; *** *P adj* < 0.001, Kruskal-Wallis test. **(F)** Flow cytometry for the mesothelial cell marker CD200 for PSCs (n=2, 3 technical replicates), fibroblasts (n=2, 2 technical replicates) and mesothelial cells (n=2, 2 technical replicates) cultured in 2D. Results show mean ± SEM. ** *P adj* < 0.01, Kruskal-Wallis test. **(G)** Flow cytometry for major histocompatibility complex class II (MHC-II) for PSCs (n=2, 3 technical replicates), fibroblasts (n=2, 2 technical replicates) and mesothelial cells (n=2, 2 technical replicates) cultured in 2D with 100 ng/mL recombinant mouse interferon gamma (IFNγ) for 48 hours. Results show mean ± SEM. * *P adj* < 0.05; ** *P adj* < 0.01, Kruskal-Wallis test. **(H)** Flow cytometry of unstained multi-stroma culture (i.e., PSCs, mesothelial cells and fibroblasts, without organoids). PSCs used are green fluorescent protein-positive (GFP^+^) and fibroblasts and mesothelial cells are TdTomato^+^. **(I-J)** Flow cytometry plot of GFP **(I)** and TdTomato **(J)** expression of PSCs (n=2), fibroblasts (n=2) and mesothelial cells (n=2) cultured in 2D. **(K)** Condition contribution to different cell subsets in murine P (n=2) and PDAC (n=2) organoid/multi-stroma co-cultures, represented as bar plot showing proportions of the different conditions in each cluster, as analysed by scRNA-seq. **(L)** Cell type contribution in P and PDAC organoid/multi-stroma co-cultures, represented as bar plots showing proportions of the different cell clusters in each condition. **(M)** Heatmap of scaled expression of cell subset specific markers of murine P and PDAC organoid/multi-stroma co-cultures as assessed by scRNA-seq. Data are scaled such that the cluster with the lowest average expression = 0 and the highest = 1 for each gene. **(N)** GSEA of the PDAC malignant signature in PDAC organoids compared to P organoids from organoid/multi-stroma co-cultures. The signature was significantly enriched in PDAC malignant cells. The PDAC malignant cell signature was obtained by comparing murine PDAC malignant cells to pancreatitis ductal cells from the snRNA-seq dataset. **(O)** Left: GSEA of the PDAC malignant signature in P organoid 1 compared to P organoid 2 cells from organoid/multi-stroma co-cultures. Right: GSEA of the pancreatitis Ductal 1-unique signature in P organoid 1 compared to P organoid 2 cells from organoid/multi-stroma co-cultures. The signature was obtained by comparing pancreatitis Ductal 1 to Ductal 2 cells from the snRNA-seq dataset of murine tissues. Both signatures were significantly enriched in ductal 1 cells. **(P)** Venn diagrams of differentially expressed curated protein-coding ortholog genes significantly (*P* adj < 0.05) upregulated in P organoids in multi-stroma co-cultures compared to PDAC organoids in multi-stroma co-cultures (hereon, ‘P organoid orthologs from multi-stroma co-cultures’), as assessed by scRNA-seq, murine pancreatitis ductal orthologs and human pancreatitis ductal orthologs. Significance of the overlap shown was defined by hypergeometric test (with denominator = 16,076 orthologs). Selected genes common to two or all datasets are shown. **(Q)** Venn diagrams of differentially expressed curated protein-coding ortholog genes significantly (*P* adj < 0.05) upregulated in PDAC organoids in multi-stroma co-cultures compared to P organoids in multi-stroma co-cultures (hereon, ‘PDAC organoid orthologs from multi-stroma co-cultures’), as assessed by scRNA-seq, murine PDAC malignant orthologs, and human PDAC malignant orthologs. Significance of the overlap shown was defined by hypergeometric test (with denominator = 16,076 orthologs). Selected genes common to two or all datasets are shown.

**Extended data Figure 7.**
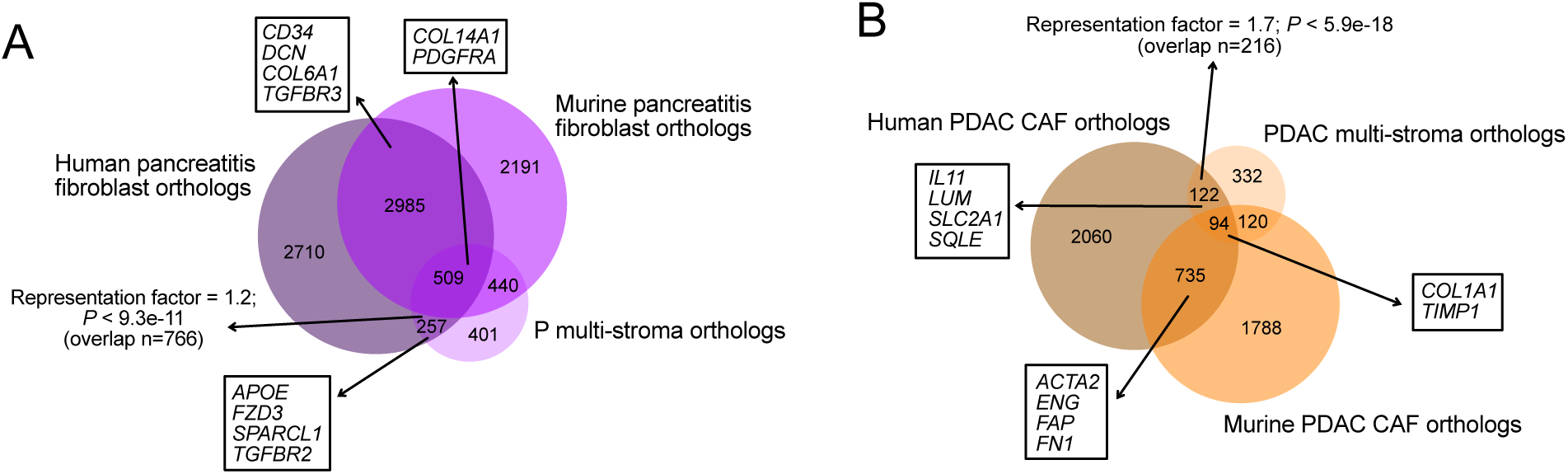
Optimised multi-stromal organoid co-cultures better mimic fibroblast heterogeneity of pancreatitis and PDAC. **(A)** Venn diagrams of differentially expressed curated protein-coding ortholog genes significantly (*P* adj < 0.05) upregulated in multi-stroma (i.e., PSCs, fibroblasts and mesothelial cells) cultured with P organoids compared to multi-stroma cultured with PDAC organoids (hereon, ‘P multi-stroma orthologs’), pancreatitis fibroblast orthologs, and human pancreatitis fibroblast orthologs, as assessed by snRNA-seq and scRNA-seq. Significance of the overlap between datasets was defined by hypergeometric test (with denominator = 16,076 orthologs). Selected genes common to two or all datasets are indicated. **(B)** Venn diagrams of differentially expressed curated protein-coding ortholog genes significantly (*P* adj < 0.05) upregulated in multi-stroma cultured with PDAC organoids compared to multi-stroma cultured with P organoids (hereon, ‘PDAC multi-stroma orthologs’), PDAC CAF orthologs, and human PDAC CAF orthologs, as assessed by snRNA-seq and scRNA-seq. Significance of the overlap between datasets was defined by hypergeometric test (with denominator = 16,076 orthologs). Selected genes common to two or all datasets are indicated.

## Supplementary Tables

**Supplementary Table 1. Single-cell RNA-sequencing of human pancreatitis and PDAC.**

**Supplementary Table 2. Single-nuclei RNA-sequencing of murine pancreatitis and PDAC.**

**Supplementary Table 3. Jaccard analyses of overlaps of differentially expressed genes from in vitro and in vivo murine pancreatitis and PDAC.**

**Supplementary Table 4. RNA-sequencing of murine normal, P and PDAC organoids in complete media.**

**Supplementary Table 5. RNA-sequencing of murine organoids sorted from P and PDAC monocultures or co-cultures with PSCs in reduced media.**

**Supplementary Table 6. RNA-sequencing of murine PSCs sorted from monocultures or co-cultures with P and PDAC organoids.**

**Supplementary Table 7. Single-cell RNA-sequencing of P and PDAC organoid/multi-stroma co-cultures.**

**Supplementary Table 8. Cross-model overlaps.**

